# Temporo-Occipital and Medial Temporal Networks Underlying Object-Location Learning

**DOI:** 10.64898/2026.02.13.705708

**Authors:** Mohamed Abdelmotaleb, Harun Kocataş, Leonardo M. Caisachana Guevara, Filip Niemann, Alireza Shahbabaie, Philipp Kuhnke, Robert Malinowski, Daria Antonenko, Gesa Hartwigsen, Marcus Meinzer, Agnes Flöel

## Abstract

Object-location memory (OLM), a fundamental component of spatial memory, is essential for everyday functioning yet declines with aging and neurodegenerative disorders. Functional neuroimaging studies have consistently implicated medial temporal lobe (MTL) structures, including the hippocampus and parahippocampal gyrus, as well as medial and lateral temporo-occipital regions, in object-location associative learning. However, the neural mechanisms and functional network interactions supporting OLM acquisition remain largely unknown. To address this gap, we examined twenty healthy adults (18–45 years) performing an object-location learning task during functional MRI. As a first main finding, task-related functional connectivity analyses revealed enhanced coupling between MTL structures, ventral visual regions, and temporo-occipital cortices during OLM learning. Secondly, stronger connectivity within this network was associated with higher learning accuracy, highlighting the behavioral relevance of coordinated cortical-MTL interactions.

Thirdly, these connectivity patterns were not static but evolved across learning stages: interregional coupling was strongest during early learning and progressively attenuated as performance stabilized, suggesting sharpening and increased efficiency of network interactions with learning. Together, our results provide mechanistic insight into OLM acquisition at the functional network level and offer an evidence-based framework for identifying target networks to enhance spatial memory through non-invasive brain stimulation.

## Introduction

Object-location memory (OLM) is a critical component of spatial memory that enables us to remember where items are located in our environment. This ability is essential for everyday functioning and is particularly vulnerable to the effects of aging—a decline that is further accelerated in neurodegenerative disorders (Lee et al., 2003; Hedden & Gabrieli, 2004; Kessels et al., 2007; Postma et al., 2008; Kessels et al., 2010). The medial temporal lobe (MTL), including the hippocampus and parahippocampal cortex, acts as a hub for binding object identity with spatial context (Hayes et al., 2004; Buffalo et al., 2006; Hannula & Ranganath, 2008; Squire & Wixted, 2011). Beyond the MTL, neuroimaging studies have consistently highlighted the conjoint engagement of a broader cortical network encompassing ventral visual-stream areas and temporo-occipital regions during object-location encoding and retrieval (Hales & Brewer, 2013; Rolls, 2024; Rolls, Zhang, et al., 2024; Abdelmotaleb et al., 2025).

Despite converging evidence for cortical-subcortical involvement in OLM, the underlying network-level interactions that support learning remain unclear. In particular, it has yet to be clarified how cortical nodes and hippocampal structures interact to facilitate successful memory formation. Investigations of visual pathways projecting to the hippocampal system have identified two complementary processing streams: (1) a ventrolateral pathway, linking the fusiform and inferior temporal cortices with the parahippocampal cortex and hippocampus, primarily associated with object-related processing; and (2) a ventromedial pathway, connecting ventromedial visual regions with the medial parahippocampus and hippocampus, associated with spatial aspects of processing (Edmund T Rolls et al., 2023; Edmund T. Rolls et al., 2023; Rolls, 2023). Furthermore, recent work contrasting task-evoked connectivity across distinct episodic memory domains—including OLM, reward-location learning, and word-pair association—has revealed domain-specific differences in functional network organization (Rolls, Zhang, et al., 2024). However, the task-evoked networks uniquely engaged by OLM relative to a matched control condition, and their relationship to individual learning success, have not been investigated yet. Addressing this gap is crucial for delineating the connectivity patterns specifically supporting object-location binding.

Characterizing these connectivity patterns is not only of theoretical importance to improve current models on the functional neuroanatomy of OLM, but also relevant for translational research. For example, identifying cortical regions that exhibit strong coupling with hippocampal memory circuits could inform network-guided targeting strategies for non-invasive brain stimulation (NIBS) interventions (Fox et al., 2014; Wang et al., 2014; Jeong et al., 2015; Nilakantan et al., 2017; Tang et al., 2023; Shahbabaie et al., 2026). Such approaches have clear clinical significance, as previous attempts to enhance OLM via NIBS have produced mixed outcomes (Flöel et al., 2012; Külzow et al., 2014; England et al., 2015; Prehn et al., 2017; Antonenko et al., 2018; Fromm et al., 2024), which may in part reflect suboptimal target selection (Meinzer et al., 2024).

Functional connectivity analyses based on generalized psychophysiological interactions (gPPI) provide a powerful framework for examining task-dependent network interactions. The gPPI approach estimates task-specific changes in interregional coupling by incorporating condition-specific regressors and their interaction terms into a connectivity model, thereby offering high sensitivity for detecting task-related modulation of functional connectivity above and beyond task-independent correlations and co-activations (McLaren et al., 2012; Masharipov et al., 2024).

Accordingly, the present study had two primary aims. First, we sought to identify functional connectivity patterns uniquely elicited during OLM relative to a closely matched non-associative control condition, using gPPI applied to a novel fMRI adaptation of a well-established OLM paradigm (Flöel et al., 2012; Külzow et al., 2014; Abdelmotaleb et al., 2025). Second, we tested whether individual differences in OLM-specific connectivity predict memory performance by correlating gPPI-derived coupling estimates with successful OLM acquisition.

We hypothesized that OLM learning would be associated with increased task-dependent functional connectivity between MTL structures and temporo-occipital regions relative to a matched control condition. Specifically, we predicted stronger coupling during learning between the hippocampus and parahippocampal cortex and ventral visual-stream regions implicated in OLM, including the fusiform gyrus, inferior and middle temporal gyri, and lateral occipital cortex (Gillis et al., 2016; Rolls, Zhang, et al., 2024; Abdelmotaleb et al., 2025). Furthermore, we hypothesized that individual differences in hippocampal–temporo–occipital connectivity strength would be positively associated with OLM performance.

In summary, this work aims to elucidate how distributed cortical–MTL networks support object-location binding and provide an empirical foundation for future network-informed neuromodulation approaches targeting this ecologically vital memory system.

## 2. Methods

This study presents findings from the preparatory phase of a larger multicenter project employing transcranial direct current stimulation (tDCS). The project is designed as a double-blind, sham-controlled, crossover study aimed at assessing the effects of tDCS on behavioral and neural responses across a range of cognitive and motor domains (for further details, see https://www.memoslap.de/de/forschung/).

### 2.1 Participants and Study Overview

A total of 20 healthy adults (aged 18–45 years) were recruited. One participant was excluded due to insufficient data quality identified through the standardized quality control pipeline (Nieto-Castanon & Whitfield-Gabrieli, 2022; Morfini et al., 2023, see Supplementary Materials, Quality Control section). The final sample comprised 19 participants (10 females; M = 24.5 years, SD = 5.2). All participants were right-handed, native German speakers, and reported no history of neurological or psychiatric disorders, substance use disorder, or other major medical conditions. Cognitive abilities were assessed using a comprehensive neuropsychological test battery covering verbal memory, visuospatial memory, and executive functions; all participants demonstrated performance within the normal range (for details, see Abdelmotaleb et al., 2025).

### 2.2 Experimental Design

Detailed experimental procedures are reported in Abdelmotaleb et al. (2025) and summarized here (Figure 1). Each participant completed two fMRI sessions, spaced at least one week apart, and performed two parallel and counterbalanced versions (A and B) of the OLM task, adapted from previous studies (Flöel et al., 2012; Prehn et al., 2017; Antonenko et al., 2018; de Sousa et al., 2020).

**Figure 1.**
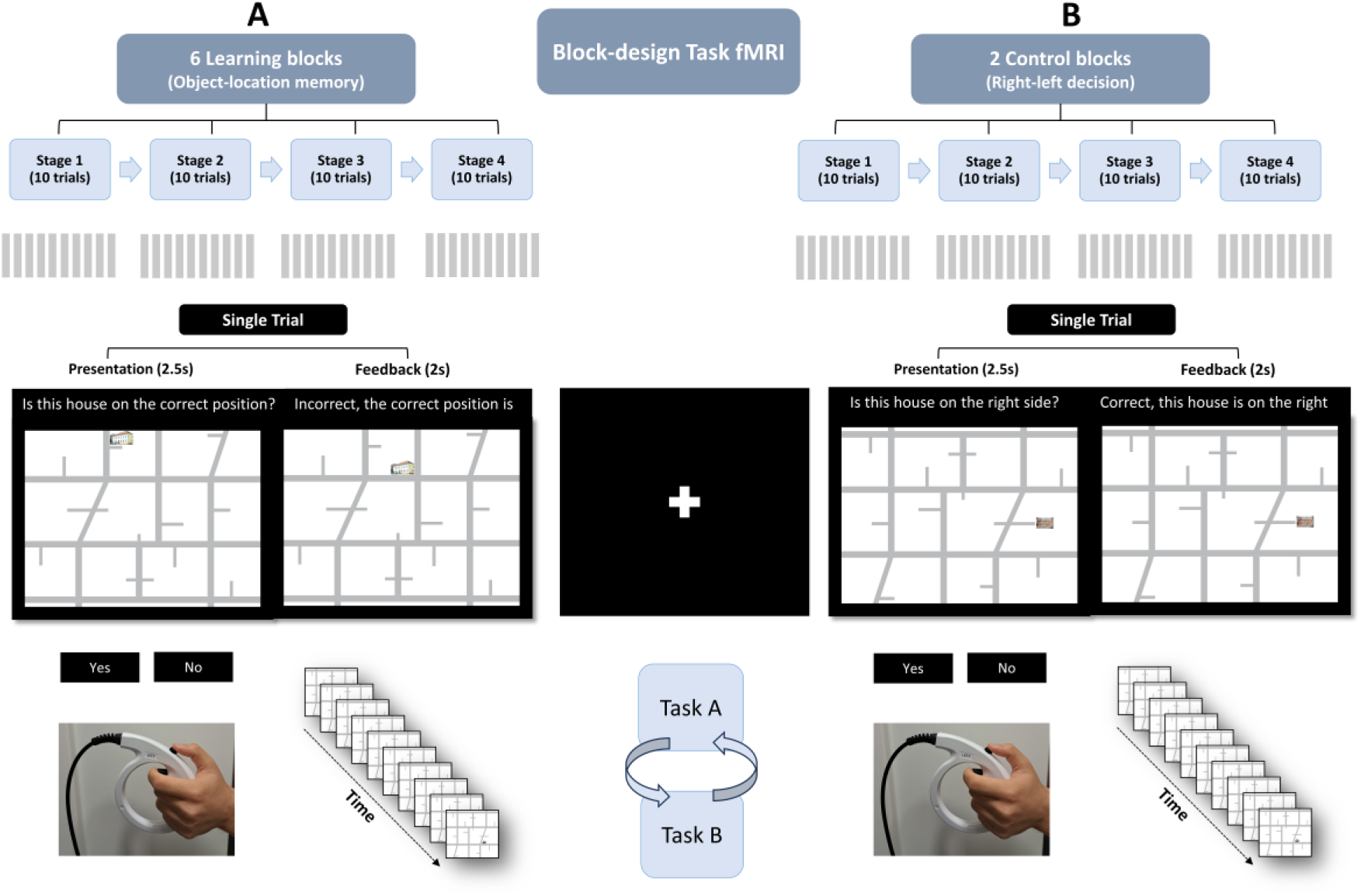
Experimental Design of the Object-Location Memory (OLM) Task. (A) fMRI block design for the learning condition, consisting of four stages per block. (B) Control blocks with the same structure but involving non-associative judgments. In the OLM task (left), participants determined whether a house was correctly positioned on a map and received feedback. In the control task (right), participants indicated whether the house appeared on the left or right side of the map, with feedback provided in the same manner. Each trial included a 2.5-second stimulus presentation and response phase, followed by 2 seconds of feedback. Two task versions (A and B) were counterbalanced across participants.

The OLM task required participants to learn associations between houses and their positions on a two-dimensional street map through repeated trials with feedback. In each trial, participants judged whether a house was correctly placed on the map and then received immediate feedback indicating both accuracy and correct location. At the beginning of each block, novel house-location pairings were introduced without prior exposure, requiring participants to make an initial guess and gradually learn through feedback. A control task, matched in structure and timing, involved left/right spatial judgments without memory demands to control for perceptual and motor components. Both the OLM and control tasks were organized into blocks, each consisting of four learning (or non-learning) stages, to minimize potential confounds such as scanner drift or fatigue (Sliwinska et al., 2017).

During each OLM block, participants viewed five unique houses presented in both correct and incorrect map locations, resulting in 10 trials per stage. Across six learning blocks per session, participants learned 30 distinct object-location associations. Control blocks followed the same structure but used 10 unique house stimuli for non-associative spatial decisions (for details, see Figure 1).

Fixation crosses (4.5 s) separated stages with longer fixation periods (18 s) interspersed between task blocks to allow the hemodynamic response function (HRF) to return to baseline. Task instructions were presented before the first session and at the start of each block using brief written prompts.

Stimuli consisted of 90 real-world house images and schematic maps divided into quadrants with randomized correct and incorrect locations. Versions A and B used distinct stimuli and rotated maps to ensure consistent difficulty across sessions. Stimuli were presented using Presentation® software (Version 20.1, Neurobehavioral Systems, Inc.) and displayed via MRI-compatible screen and mirror system. Performance data were collected using MRI-compatible response grips (Nordic Neuro, Norway).

To establish a practical workflow for future combined tDCS-fMRI experiments, sham tDCS was administered during functional imaging sessions (for procedural details, see Niemann et al., 2024 and Abdelmotaleb et al., 2025). As a preliminary cortical target, the right lateral occipito-temporal cortex (ROTC) was selected based on previous neuroimaging evidence (Gillis et al., 2016; Thielscher et al., 2026).

### 2.3 fMRI Data

MRI data were acquired at University Medicine Greifswald using a 3.0 Tesla Siemens MAGNETOM Vida scanner (Siemens Healthineers, Germany) equipped with a 64-channel head-neck coil.

Functional images were obtained with a multiband gradient-echo-planar imaging (EPI) sequence (TR = 1,000 ms; TE = 30.8 ms; flip angle = 60°; field of view = 220 × 220 mm; acquisition matrix = 110 × 110; 72 axial slices; no inter-slice gap; voxel size = 2 × 2 × 2 mm; multiband acceleration factor = 6; GRAPPA within-slice acceleration factor = 2).

High-resolution structural images were collected using a T1-weighted MPRAGE sequence (TR = 2,700 ms; TE = 3.7 ms; TI = 1,090 ms; flip angle = 9°; field of view = 259.2 × 259.2 mm; acquisition matrix = 288 × 288; 224 sagittal slices; voxel size = 0.9 × 0.9 × 0.9 mm).

Functional and anatomical MRI data were preprocessed using the CONN toolbox (Whitfield-Gabrieli & Nieto-Castanon, 2012; Nieto-Castanon & Whitfield-Gabrieli, 2022; RRID: SCR_009550; release 22.v2407) in combination with SPM12 (UCL, London, UK; version 12.7771; RRID: SCR_007037) in MATLAB (The MathWorks Inc., Natick, MA, USA; R2021b). Preprocessing followed CONN’s modular pipeline (Nieto-Castanon, 2020) including: realignment with correction for susceptibility distortion interactions, slice timing correction, outlier detection (outliers were defined as framewise displacement > 0.9 mm or global blood oxygenation level dependent (BOLD) signal > 5 SD; Power et al., 2014; Whitfield-Gabrieli et al., 2011), anatomical segmentation, normalization to MNI space via unified segmentation of the coregistered anatomical image (resampled to 2-mm isotropic voxels), and spatial smoothing using a 6-mm full-width at half-maximum (FWHM) Gaussian kernel.

Functional data were denoised using a standard CONN toolbox pipeline (Nieto-Castanon, 2020), which involved regression of multiple sources of nuisance variance. These included five aCompCor components each from white matter and CSF time series (Behzadi et al., 2007; Chai et al., 2012), six motion parameters and their first-order derivatives (12 regressors in total; Friston et al., 1996), up to 65 outlier volumes identified during preprocessing (Power et al., 2014), session and task effects along with their first-order derivatives, and a constant term for each functional run (Whitfield-Gabrieli & Nieto-Castanon, 2012; Fair et al., 2007; Nieto-Castanon, 2020). Following nuisance regression, the BOLD time series was temporally band-pass filtered (0.008-0.09 Hz) to retain physiologically relevant low-frequency signal fluctuations (Hallquist et al., 2013).

Given that motion and other sources of noise can substantially compromise fMRI data quality, particularly in functional connectivity analyses (Power et al., 2012), we implemented a rigorous quality control (QC) protocol across raw data, preprocessing, and denoising stages, following established pipelines (Nieto-Castanon & Whitfield-Gabrieli, 2022; Morfini et al., 2023). Details are in the supplementary materials (Quality control section and Figure S1).

### 2.4 Functional Connectivity Analysis

Task-dependent functional connectivity was examined using generalized psychophysiological interaction (gPPI) analyses for the two experimental conditions (learning and control).

Psychophysiological interaction (PPI) analysis is commonly used to identify brain regions whose connectivity with a given seed region varies as a function of experimental context (Friston et al., 1997). The gPPI framework models condition-specific connectivity effects and is particularly well suited for block-design paradigms, offering improved sensitivity and specificity relative to traditional PPI approaches (McLaren et al., 2012; Kuhnke et al., 2021; Masharipov et al., 2024).

For each seed-target pair, a gPPI model was specified, incorporating three components: the seed region’s BOLD time series (physiological regressor), condition-specific boxcar regressors convolved with the canonical HRF (psychological regressors), and their interactions (psychophysiological terms). Connectivity analyses were conducted using both region of interest-to-region of interest (ROI-to-ROI) and seed-to-voxel approaches.

#### ROI-to-ROI Connectivity Analyses

Pairwise functional connectivity was quantified across 132 anatomically defined ROIs from the Harvard-Oxford cortical and subcortical probabilistic atlas (https://fsl.fmrib.ox.ac.uk/fsl/fslwiki/; Frazier et al., 2005; Makris et al., 2006; Desikan et al., 2006; Goldstein et al., 2007). Connectivity matrices were estimated using CONN, and the resulting matrices were subsequently imported into Python for visualization using Nilearn (Nilearn contributors, n.d.; Abraham et al., 2014, RRID:SCR_001362) and Nichord (Bogdan et al., 2023).

For clarity of presentation, the Results section displays only connectivity edges (i.e., pairwise functional connections between ROIs) involving OLM-relevant regions that have been consistently implicated in object-location learning, including the hippocampus, parahippocampal gyrus, fusiform gyri, temporo-occipital cortex, and lateral occipital cortex (Postma et al., 2008; Gillis et al., 2016; Rolls, Zhang, et al., 2024). Complete ROI-to-ROI connectivity results for all atlas-defined ROIs are provided in the Supplementary Materials.

#### Seed-to-Voxel Analyses

Additional seed-to-voxel analyses were subsequently performed to complement and validate the ROI-to-ROI findings and to characterize whole-brain functional connectivity patterns associated with each OLM-relevant seed region (Supplementary Materials for details).

#### Group-Level Inference

We conducted group-level one-sample t-tests on the subject-level contrast maps for learning > control connectivity. Group statistical maps were thresholded at a voxel-wise p < 0.001, with cluster-wise family-wise error correction (FWE) at p < 0.05.

### 2.5 Brain-Behavior Correlations

To examine whether task-related functional connectivity predicted performance on the OLM task, we correlated ROI-to-ROI gPPI connectivity estimates from the learning condition with behavioral accuracy, our primary outcome measure. Analyses were hypothesis-driven and restricted to connections within an a priori OLM network, as defined above. Connectivity estimates consisted of standardized subject-level beta coefficients derived from the ROI-to-ROI gPPI models. This ROI-to-ROI framework provided edge-wise connectivity measures, enabling direct statistical comparisons across participants and facilitating correlation with behavioral performance.

Behavioral performance was indexed by accuracy. For across-subject analyses, a single performance metric per participant was computed as the mean accuracy across all learning stages. For stage-specific analyses, accuracy was calculated separately for each learning stage (Stages 1-4) for each participant (n = 19).

Connectivity–behavior relationships were examined at three complementary analytical levels:

1. Across-subject (mean-level) correlations: To assess whether individuals with stronger overall task-related connectivity performed better or worse, we correlated mean gPPI connectivity values for each ROI pair within the a priori network with participants’ mean accuracy scores (Pearson’s r).
2. Stage-specific across-subject correlations: To determine whether connectivity tracked performance at each learning stage, we estimated gPPI connectivity using separate regressors for stages 1–4. Connectivity values were then correlated with accuracy across participants at each stage (yielding four Pearson correlations per ROI pair). This approach preserved the stage structure while testing between-subject variation in performance.
3. Within-subject (stage-wise) correlations: To quantify the relationship between connectivity and accuracy across learning stages within individuals, we used repeated-measures correlation (rmcorr; Bakdash & Marusich, 2017).

Rmcorr estimates a common slope between connectivity and accuracy while allowing for participant-specific intercepts, thereby controlling for between-subject differences and isolating within-subject associations across Stages 1–4. For each ROI pair, we report the rmcorr coefficient (r), associated p-value, and the estimated common slope.

All connectivity–behavior correlation analyses across analytical levels were implemented in Python (version 3.11.5) using standard libraries, including NumPy, pandas, SciPy (Virtanen et al., 2020), Pingouin (Vallat, 2018) and statsmodels (Seabold & Perktold, 2010). Significance was initially evaluated at p < 0.05 (uncorrected) for a cross-subject analyses (Levels 1 and 2). For the repeated-measures correlation analyses (Level 3), we applied Holm’s step-down procedure (α=0.05) to control family-wise error for all edges in that hypothesis set.

## Results

### 3.1 OLM Functional Connectivity

To provide context for the subsequent connectivity analyses, we first summarize the task-evoked activation patterns observed during the Object-Location Memory (OLM) task, based on the same dataset (see Abdelmotaleb et al., 2025, for full details). During OLM, significant activation was observed bilaterally in temporo-occipital regions, including the fusiform cortex, hippocampus, parahippocampal gyrus, inferior temporo-occipital gyrus, and lateral occipital cortex. Additional engagement was identified in subcortical regions, precentral and postcentral gyri, orbitofrontal cortex, and precuneus. By contrast, the control > learning contrast showed stronger activity in occipital and lingual visual cortices. An adapted overview of these activation patterns is presented in Figure 2 (for comprehensive statistical results, see Abdelmotaleb et al., 2025).

**Figure 2.**
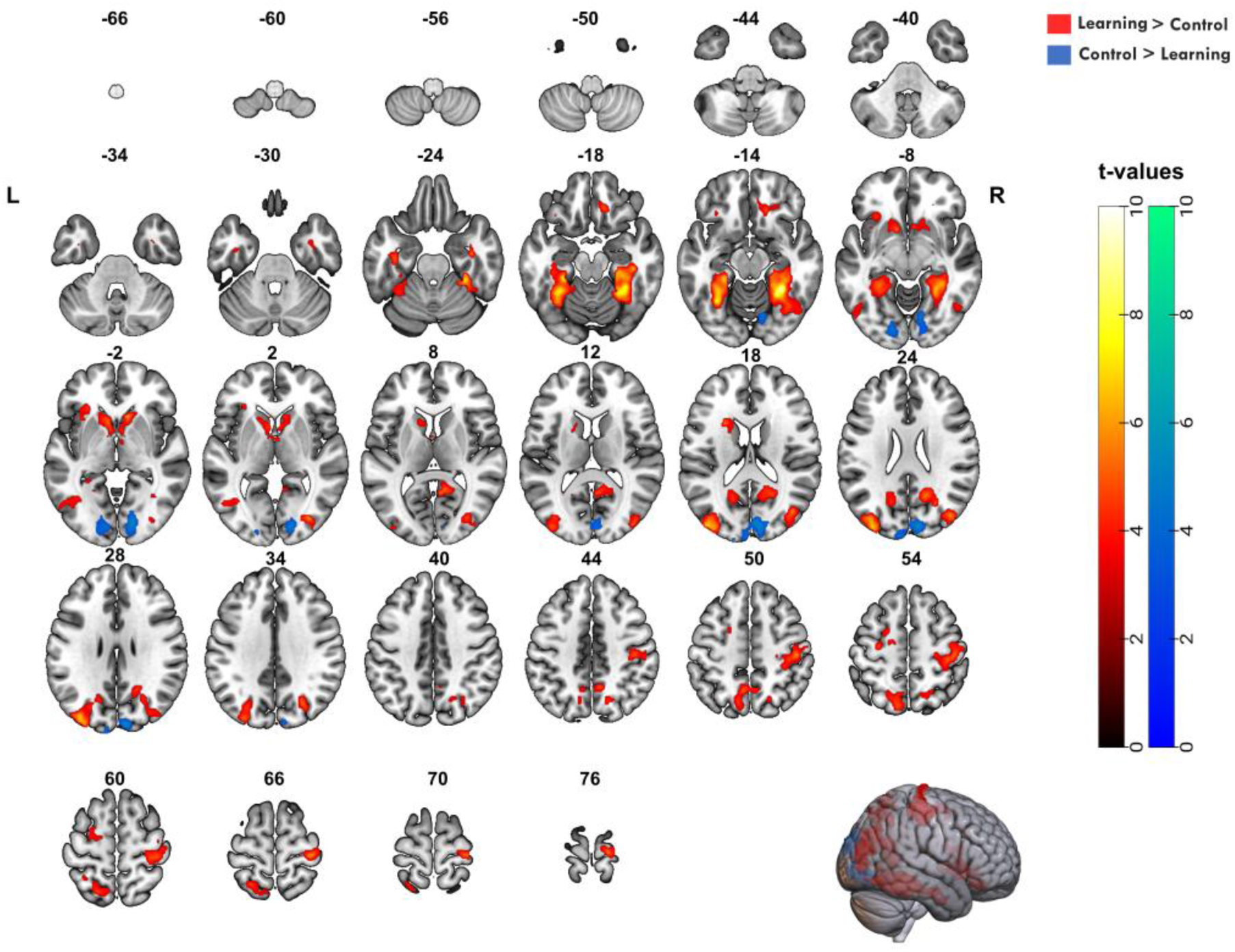
Brain activation patterns during the learning and control tasks. Group-level statistical parametric maps projected onto an MNI template. Warm-colored clusters indicate regions with greater activation for the learning > control contrast, whereas cool-colored clusters denote areas with stronger activation for the control > learning contrast. Analyses were conducted across two task sessions, with results thresholded at a voxel-wise p < 0.001 and a cluster-wise p < 0.05 FWE-corrected. The color bar indicates t-values, and activations are displayed in axial orientation. Adapted from Abdelmotaleb et al., 2025.

#### 3.1.1 ROI-to-ROI Analysis

ROI-to-ROI functional connectivity analyses were conducted to characterize pairwise interactions among predefined regions of interest (ROIs) for the learning > control contrast. Connectivity was quantified between all ROI pairs defined by the Harvard-Oxford atlas.

We first estimated the full connectivity matrix comprising 132 atlas-based ROIs. For visualization and interpretability, we subsequently focused on connections involving OLM-relevant regions, including the hippocampi, parahippocampal gyri, fusiform gyri, and temporo-occipital/lateral occipital cortices. These connections are illustrated as a connectome highlighting hippocampal and parahippocampal interactions (Figure 3A) and as a heatmap of significant edges (Figure 3B).

**Figure 3.**
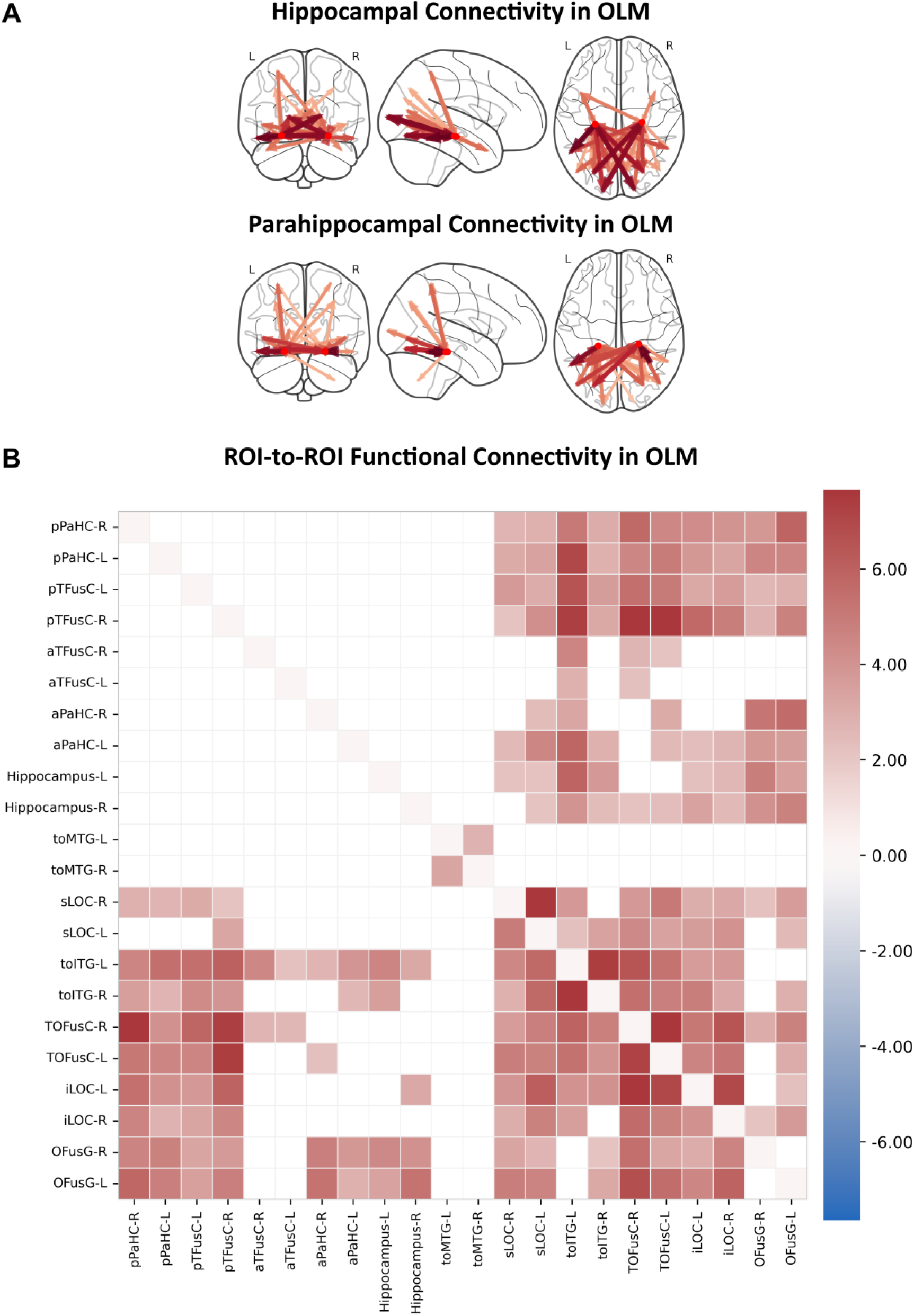
Task-based ROI-to-ROI functional connectivity during object-location learning. (A) Connectome plots illustrating significant ROI-to-ROI functional connectivity differences between the object-location learning and control conditions. Illustrations depict connectivity patterns involving the bilateral hippocampi and the parahippocampal cortices. (B) Group-level ROI-to-ROI connectivity matrix (19 participants) derived from gPPI analysis for the contrast learning > control, based on a predefined set relevant to OLM from Harvard-Oxford-ROIs. The color scale indicates t-values, with warm colors denoting stronger connectivity during learning relative to control. No significant negative correlations were found.

The hippocampus and parahippocampal gyri, key hubs for OLM, exhibited widespread connectivity during learning. Both hippocampi showed enhanced coupling with bilateral fusiform gyri and with the temporo-occipital portions of the inferior temporal gyrus. The right parahippocampal gyrus showed robust connectivity with bilateral temporo-occipital fusiform, bilateral occipital fusiform, bilateral lateral occipital cortices, and bilateral temporo-occipital inferior temporal gyri. The left parahippocampal gyrus demonstrated a similar pattern, connecting with the left temporo-occipital inferior temporal gyrus, left lateral occipital cortex, bilateral temporo-occipital fusiform, and bilateral occipital fusiform.

The temporo-occipital cortices, encompassing bilateral middle and inferior temporal gyri, also showed strong intra- and interhemispheric coupling. The middle and inferior temporo-occipital regions were significantly connected within each hemisphere and across hemispheres. The bilateral inferior temporal gyri exhibited particularly strong connectivity. Additionally, the right inferior temporo-occipital cortex connected with bilateral fusiform gyri and the left lateral occipital cortex, while the left inferior temporo-occipital cortex demonstrated reciprocal connectivity with bilateral fusiform, bilateral lateral occipital cortices, as well as with hippocampal and parahippocampal regions.

All significant edges, along with corresponding t-statistics and p-values, are reported in supplementary materials (Table S17). A comprehensive visualization of the full atlas ROI-to-ROI connectivity matrix is provided in Supplementary Figure S4.

To confirm the ROI-to-ROI findings and to characterize whole-brain connectivity patterns within the OLM network, we conducted additional seed-based functional connectivity analyses. Seed regions were selected based on their involvement in learning-related activity and included the hippocampus, parahippocampal gyrus, temporo-occipital cortex, and lateral occipital cortex, as defined by the Harvard-Oxford atlas. Whole-brain connectivity maps for each seed in the learning > control contrast are shown in Supplementary Figure S2. Across seeds, OLM learning was associated with a robust and coherent connectivity pattern linking MTL structures with ventral visual and occipito-temporal cortices, a pattern that was largely absent during the control condition.

Full statistical details for all seed-based analyses are provided in Supplementary Tables S1-S10.

### 3.2. Brain–Behavior Relationships

#### 3.2.1 Behavioral Data Analysis

As reported in detail elsewhere (Abdelmotaleb et al., 2025), during OLM learning, participants demonstrated robust learning across the four stages of feedback-based learning. Accuracy increased and reaction times (RTs) decreased as learning progressed (For details, see Supplementary Figure S7). Generalized linear mixed-model (GLMM) analyses revealed significant main effects of stage and task type, as well as their interaction. Accuracy was initially higher in the control condition (Stages 1–2), converged with the learning condition by Stage 3, and was surpassed by the learning condition at Stage 4, confirming successful acquisition of object-location associations. RTs decreased across stages but remained consistently longer for the learning condition, reflecting the higher cognitive demands of the memory task despite progressive efficiency gains.

#### 3.2.2 Connectivity-Behavior Correlations

We assessed the relationship between task-related functional connectivity and OLM performance using three levels of analysis derived from ROI-to-ROI gPPI estimates in the learning condition.

First, ROI-to-ROI gPPI connectivity values during the learning condition were correlated with mean accuracy across participants. Across the predefined subset of ROIs, several significant connectivity-behavior correlations emerged (Figure 4). Positive correlations were primarily observed between medial temporal lobe structures and ventral visual cortices. Notably, stronger connectivity between the right inferior temporal gyrus (temporo-occipital) and the right posterior parahippocampal gyrus (r = 0.52, p = 0.024), the left inferior temporal gyrus (temporo-occipital) and the right posterior parahippocampal gyrus (r = 0.53, p = 0.018), and between left and right temporal-occipital cortices (ITG; r= 0.50, p=0.031) was associated with higher learning accuracy. Additional positive connectivity-behavior associations were observed between the right temporal occipital fusiform gyrus and the left inferior temporal gyrus (temporo-occipital) (r = 0.52, p = 0.022), and between the left hippocampus and the right occipital fusiform gyrus (r = 0.49, p = 0.035). This highlights that between-subject differences in connectivity within the OLM-relevant network predict OLM performance.

**Figure 4.**
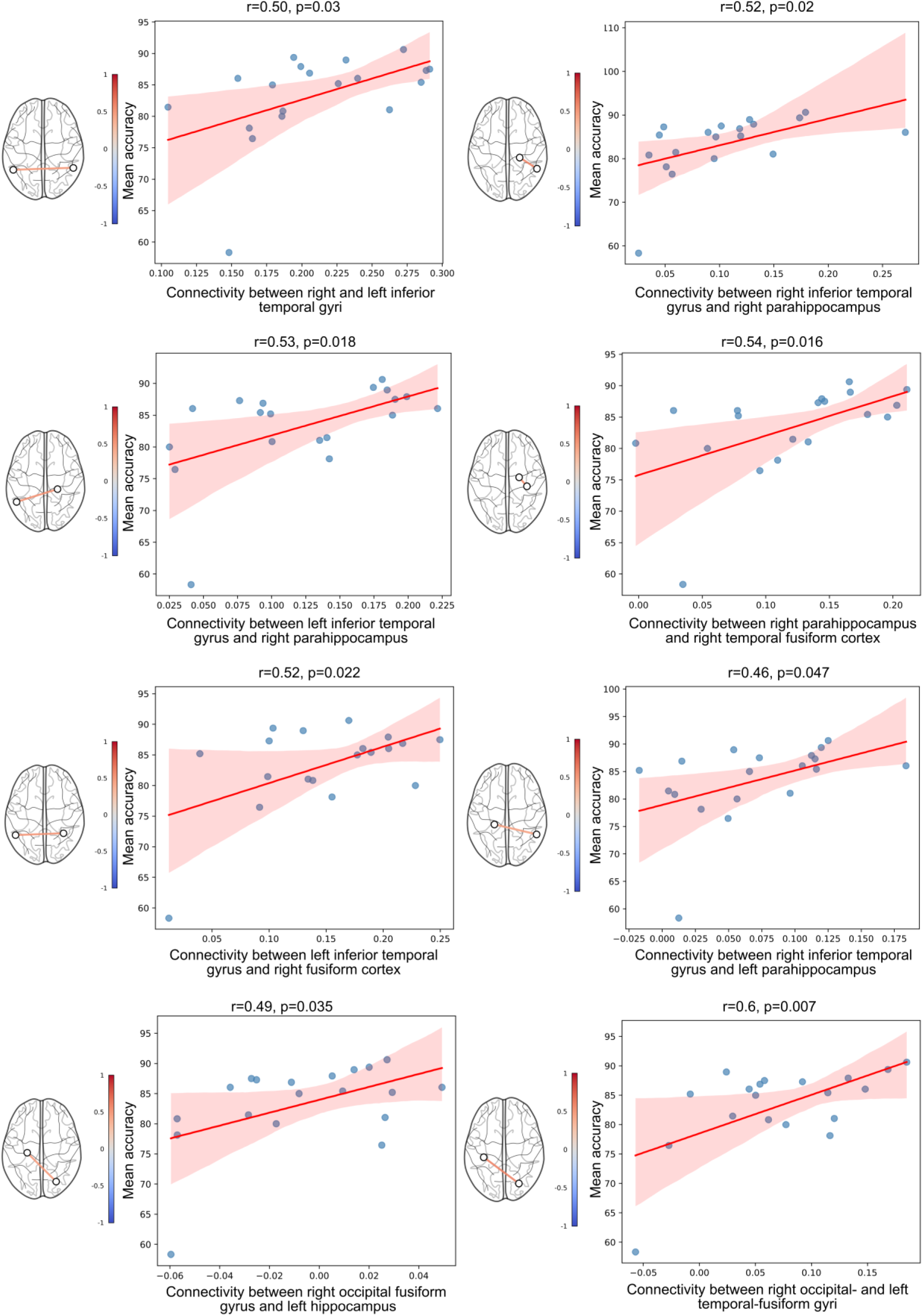
Connectivity-behavioral correlations for ROI-to-ROI gPPI connectivity during object-location learning. Scatterplots show correlations between gPPI connectivity beta values and mean learning accuracy for selected ROI pairs identified from whole-brain gPPI connectivity analysis using the Harvard-Oxford atlas. ROI pairs shown were chosen from the larger connectivity matrix based on their relevance to OLM, including connections involving bilateral hippocampi, parahippocampal gyri, fusiform gyri, and temporo-occipital portions of the middle and inferior temporal gyri. Each plot reports the correlation coefficient (r) and p-value, with a regression line and 95% confidence interval. Positive associations were primarily observed between medial temporal lobe structures and ventral visual cortices. Adjacent to each scatterplot, a brain network visualization highlights the corresponding ROI pair showing a positive connectivity–behavior relationship during learning.

Second, we examined stage-wise ROI-to-ROI gPPI connectivity-performance relationships by correlating connectivity estimates with accuracy across participants for the selected edges. Within the OLM network, significant associations were predominantly positive, such that stronger functional connectivity at a given learning stage was associated with higher performance (Supplementary Figures S5 and S6). Notably, the earliest learning stages exhibited the highest density of significant connectivity-accuracy associations within OLM-relevant ROIs, consistent with enhanced network engagement during early phases of learning (see Supplementary Figure S6 for details).

Lastly, to examine within-subject changes in connectivity across learning stages, we applied repeated-measures correlation (rmcorr). In contrast to between-subject effects, within-subject correlations in the OLM-relevant network were predominantly negative across stages (Figure 5). This pattern indicates that as participants became more proficient, connectivity within the OLM network decreased, consistent with increasing network efficiency or functional specialization over learning. Figure 5 depicts the within- vs between-subjects reversal paradox; connectivity relates positively to performance at a given stage, yet within individuals, connectivity progressively declines as learning consolidates, potentially reflecting increasing network efficiency and reduced task demands as performance approaches ceiling.

**Figure 5.**
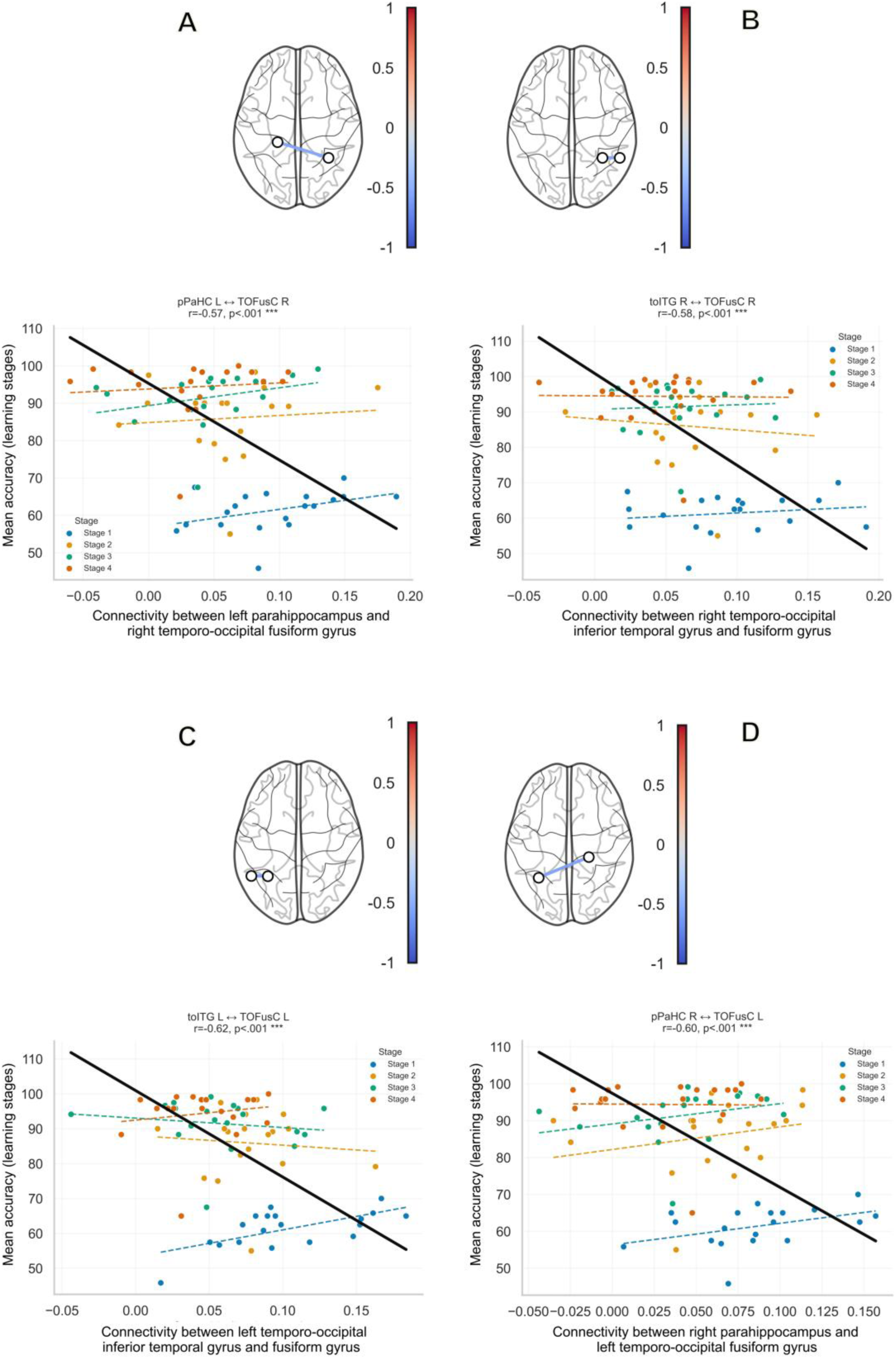
Repeated measures correlations (rmcorr) for ROI-to-ROI gPPI connectivity with learning across stages during object-location learning. Scatterplots show correlations between connectivity strength and learning accuracy across four stages of learning for selected ROI pairs identified from whole-brain gPPI connectivity analysis using the Harvard-Oxford atlas. Significant negative correlations between connectivity estimates and behavioral accuracy across learning stages in relevant OLM ROIs, including A. Parahippocampus, posteriort left to temporal-occipital fusiform gyrus right, B. Temporal-occipital fusiform gyrus right to temporooccipital inferior temporal cortex right, C. Temporooccipital inferior temporal cortex left to temporal-occipital fusiform left, D. Temporal-occipital fusiform left to parahippocampus posterior right. For each selected edge (A-D), points are colored by stage (1–4). The solid black line depicts the common rmcorr fit; dashed colored lines show stage-specific across-subject trends. Above each scatterplot, a connectome highlights the corresponding ROI pair. Reported r and p values refer to the rmcorr analysis (family-wise error corrected).

## Discussion

The present study characterized the complex functional networks supporting object-location memory (OLM) in healthy adults and identified cortical nodes with coupled medial temporal lobe (MTL) memory structures during feedback-based OLM learning. Using a gPPI functional connectivity framework, including ROI-to-ROI and seed-to-voxel analyses, together with a validated OLM paradigm, we demonstrated that OLM engaged a distributed, yet interconnected cortico-MTL network that was distinct from the network recruited by a structurally matched control task not requiring learning. This OLM network included the hippocampus, parahippocampal cortex, fusiform gyrus, temporo-occipital cortices, and lateral occipital cortices. Importantly, functional coupling within this network was behaviorally relevant: stronger connectivity between MTL structures and ventral visual cortices was associated with superior learning accuracy across individuals. Brain–behavior relationships were also dynamic. While individuals with overall stronger coupling performed better overall, within-person connectivity declined across stages of learning, consistent with increasing neural efficiency and reduced reliance on widespread inter-regional coordination as associations became consolidated and performance approached ceiling. Together, these results point to a core cortico-hippocampal network that supports associative binding of objects to locations and contributes to individual differences in OLM performance. Our results also inform potential network-based targeting approaches for enhancing OLM by non-invasive brain stimulation (NIBS).

### 4.1 Cortical–MTL Memory Network for OLM

Across ROI-to-ROI and seed-to-voxel analyses, OLM learning relative to a structurally matched control task reliably increased functional coupling between MTL structures and ventral visual cortices. Specifically, hippocampal and parahippocampal regions showed stronger interactions with occipito-temporal areas, including fusiform, inferior/middle temporal cortices and lateral occipital cortex, regions implicated in high-level visual representations (Petersson et al., 2001; Grill-Spector et al., 2006; Rolls, 2021; Rolls, Yan, et al., 2024). This pattern aligns with established models of episodic memory in which the hippocampal system integrates object identity with spatial and contextual information (Davachi, 2006; Montaldi & Mayes, 2010; Postma et al., 2004, 2008), and extends prior univariate activation findings by demonstrating task-dependent coupling among these components during feedback-driven learning (Abdelmotaleb et al., 2025; Rolls et al., 2024).

At the systems level, the present connectivity profile is consistent with recent models proposing that hippocampal episodic binding integrates both object (“what”) and viewed spatial or contextual (“where”) information conveyed predominantly via ventral temporal pathways (Rolls, 2024), rather than relying primarily on dorsal parietal routes emphasized in classical visual processing frameworks (Ungerleider & Haxby, 1994). Within this framework, the ventrolateral visual stream conveys high-level object representations through inferior temporal cortex and parahippocampal regions, whereas a ventromedial pathway involving medial visual stream areas, including fusiform and parahippocampal cortices, provides information about the currently viewed location within a scene (Edmund T Rolls et al., 2023; Edmund T. Rolls et al., 2023). These inputs converge in the hippocampus to form object-in-place episodic representations (Rolls, 2023; Rolls & Treves, 2024). In line with this account, we observed strong coupling between hippocampal/parahippocampal structures and ventral occipito-temporal regions during learning, with comparatively weaker parietal involvement.

Although activation studies of OLM often show a right-lateralized pattern (Rolls, Zhang, et al., 2024; Abdelmotaleb et al., 2025), functional coupling in the present study was largely bilateral and symmetric. This dissociation is not unexpected, as univariate activation and task-modulated connectivity index different aspects of neural processing and do not necessarily align (Satake et al., 2024). Rather than strict hemispheric dominance, OLM appears to rely on distributed bilateral networks, with right temporal regions supporting spatial computations and left homologues contributing complementary processes such as object identity or verbal labelling, coordinated through interhemispheric interactions (Postma et al., 2008; Bellgowan et al., 2009; Karolis et al., 2019; De Biase et al., 2025).

Contrasting the learning condition with a closely matched control task was crucial for isolating OLM-specific interactions. During the control condition, parahippocampal cortex and lateral occipital cortex instead showed increased connectivity with cingulate, paracingulate, supramarginal, opercular, and sensorimotor regions—networks typically associated with attentional control, salience detection, and motor processing (Vossel et al., 2014; Sadaghiani & D’Esposito, 2015). Notably, these connections did not involve the hippocampal-occipito-temporal coupling observed during learning. This dissociation indicates that the cortical–MTL interactions identified here are specific to the associative demands of object-location binding rather than reflecting generic visuospatial processing or visual stimulation.

### 4.2 Behavioral relevance of functional connectivity patterns

A central goal of the study was to evaluate whether task-evoked changes in connectivity within the OLM network have behavioral relevance. Across individuals and learning stages, higher OLM accuracy was consistently associated with stronger coupling among OLM key network nodes. This enhanced integration along the ventral visual–MTL axis confirms that interactions among episodic memory hubs are critical for successful memory formation and retrieval (Tambini et al., 2010; Schedlbauer et al., 2014). The observed correlations support the functional specialization of these regions and suggest that interindividual variability in learning success, here for the OLM task, may be at least partly due to interindividual differences in connectivity (Passiatore et al., 2025). These findings further imply that modulating connectivity within this network may facilitate the encoding of new object-location associations.

Importantly, positive brain–behavior relationships were mainly evident in early learning stages when participants were actively using feedback to establish OLM traces. Stage-wise, across-subject analyses revealed that individuals with stronger coupling within OLM-relevant edges at a given stage tended to perform better, indicating that functional connectivity may actively support the rapid acquisition of new associations. These results align with emerging evidence that dynamic reconfiguration of large-scale memory networks contributes to individual differences in learning efficiency (Shi et al., 2024).

### 4.3 Learning-related reconfiguration: efficiency and specialization

Although stronger coupling predicted better performance across participants, within-subject analyses revealed the opposite direction: as a given participant’s accuracy improved from learning stage 1 to stage 4, connectivity within the OLM network tended to decrease. We interpret this apparent paradox as evidence for efficient learning-related neural network reconfiguration. This pattern is similar to fMRI activations’ decay across stages of learning reported in both object-location and novel word learning (Abdelmotaleb et al., 2025; Kocataş et al., 2025) or decreased activity associated with better performance during tDCS (Meinzer et al., 2012; Fiori et al., 2018).

One plausible account is the efficiency hypothesis. Early in learning, participants must integrate visual identity, spatial coordinates, and feedback signals on a trial-by-trial basis. This likely requires strong communication between the ventral visual cortex, the parahippocampal cortex, and the hippocampus to form and update associations. After successful formation of OLM, retrieval may shift toward more automatic pattern completion with reduced need for widespread interregional coupling (Wander et al., 2013). In other words, encoding and feed-back learning are “network demanding”, whereas stabilized retrieval is “network efficient.” Similar decreases in cortical-hippocampal coupling with practice have been described in associative memory and category learning, where initially distributed networks give way to more focal or streamlined patterns as representations crystallize (Tambini et al., 2010; Wander et al., 2013).

A second, related interpretation pertains to functional specialization. High early coupling may reflect the joint engagement of multiple learning components: object features, location quadrants, and response contingencies. Through learning, the network may eliminate inefficient representations and stabilize into a dominant coding scheme, thereby diminishing reliance on broad cross-regional integration (Staresina et al., 2016; Sučević & Schapiro, 2023). Under this view, reduced coupling does not signal disengagement but rather sharpening.

Taken together, the across-subject and within-subject analyses suggest that strong cortico-hippocampal coupling is beneficial, but primarily at the moment when associations are still being formed and refined. Once learning has succeeded and associations are established, the same level of large-scale integration is no longer required. Collectively, these findings emphasize the value of considering both variability in overall learning across participants and within-subject variability in the trajectory across the different learning stages.

### 4.4 Toward connectivity-guided non-invasive brain stimulation

One aim of this study was to identify cortical regions that are functionally coupled with hippocampal memory circuitry during OLM formation to inform future stimulation protocols. Conventional NIBS methods, specifically transcranial magnetic stimulation (TMS) and transcranial direct current stimulation (tDCS), cannot directly stimulate the hippocampus or parahippocampal gyrus due to fundamental limitations in reaching deeper brain structures (Thair et al., 2017; Liu et al., 2022; Siebner et al., 2022). However, a growing body of evidence demonstrates that stimulating cortical nodes with strong coupling with the MTL can modulate hippocampal activity and enhance associative memory (Tambini et al., 2010; Wang et al., 2014; Nilakantan et al., 2017; Warren et al., 2019; Ester-Nacke et al., 2024). A recent meta-analysis showed that non-invasive brain stimulation directed at hippocampal-related networks reliably improves episodic memory performance across a broad range of study designs (Goicoechea et al., 2025).

Our findings highlight inferior and middle temporal cortices at the temporo-occipital junction as promising cortical access points for OLM enhancement. These regions (i) exhibited robust coupling with the hippocampus, parahippocampus, and other OLM-relevant areas; (ii) contributed to behaviorally relevant edges predicting learning accuracy; and (iii) are anatomically accessible to conventional non-invasive stimulation (Ezzyat et al., 2018; Kuhnke et al., 2025; Thielscher et al., 2026). Moreover, these regions were primarily activated during OLM and are known to connect object-specific visual codes and spatial binding processes (Rolls, Yan, et al., 2024; Abdelmotaleb et al., 2025).

Methodologically, the present study leveraged a gPPI framework to characterize task-dependent functional connectivity during object-location learning. Compared with standard PPI approaches, gPPI offers improved sensitivity and specificity in multi-condition designs by modeling condition-specific interaction terms (McLaren et al., 2012), an advantage that has been supported with high sensitivity in contrasting connectivity between task states in block designs (Masharipov et al., 2024).

Its prior successful application in associative memory research (Caviezel et al., 2020) makes it well-suited for isolating OLM-specific coupling patterns. An additional strength was the use of task-evoked, rather than resting-state, connectivity. Although resting-state networks often approximate task-related configurations and have been widely applied in memory research (Cole et al., 2014; Ritchey et al., 2014; Jeong et al., 2015; Persson et al., 2018), task-evoked connectivity is better suited to capture process-specific interactions uniquely engaged during learning (Lynch et al., 2018; Cole et al., 2021; Tuominen et al., 2023). Accordingly, our task-control design enabled the isolation of connectivity changes directly related to object-location binding.

The study also has limitations. The sample size was modest (n = 19), which may constrain statistical power, particularly for connectivity–behavior associations. The two-session design improved the stability of within-subject estimates. We further mitigated this limitation by restricting hypothesis-driven analyses to an a priori OLM-relevant network and by employing repeated-measures correlation to enhance sensitivity to within-subject effects. Nevertheless, some effects did not survive stringent correction for multiple comparisons and replication in larger samples is recommended.

### 4.5 Conclusion

This study advances our mechanistic understanding of OLM in three key aspects. First, it delineates the functional architecture through which MTL structures interact with cortical systems to bind object identity to spatial context. Second, it demonstrates that functional connectivity within this network is behaviorally relevant, as stronger network coupling is associated with greater learning success. Third, it identifies cortical, stimulation-accessible nodes that are tightly linked to hippocampal memory processes during OLM formation. By bridging systems-level memory neuroscience with network-guided intervention principles, this work lays empirical groundwork for targeted modulation of OLM using NIBS approaches.

## Ethics statement

The study was conducted at University Medicine Greifswald and received ethical approval from the institution’s medical ethics committee (approval number BB015/22). All procedures adhered to the Declaration of Helsinki, and written informed consent was obtained from all participants prior to enrolment.

## Data Availability Statement

The research protocol was pre-registered on the Open Science Framework (OSF), and the full registration details are publicly accessible at https://osf.io/t37u2. The fMRI data as well as codes will be made openly available via the EBRAINS platform, in accordance with FAIR data-sharing principles.

## Acknowledgments

This research was funded by the German Research Foundation (DFG) (project grants: ResearchUnit5429/1 [467143400]; CRC1315-B03 [327654276]; FL379/22-1; FL379/26-1; FL379/34-1; FL379/35-1; FL379/37- 1; ME3161/3-1; ME3161/5-1; ME3161/6-1; AN1103/3-1; AN1103/6-1 [539593253] to DA, AN 1103/4-1 [497919823] to DA). We are grateful to our research coordinators and student assistants for their help with scheduling and testing.

## References

Abdelmotaleb, M., Niemann, F., Kocataş, H., Caisachana Guevara, L. M., Shahbabaie, A., Malinowski, R., Riemann, S., Fromm, A. E., Hayek, D., Antonenko, D., Meinzer, M., & Flöel, A. (2025). Identification of Reliable Target Brain Regions for Enhancing Object-Location Memory by Brain Stimulation. Brain and Behavior, 15(7), e70658. 10.1002/brb3.70658

Abraham, A., Pedregosa, F., Eickenberg, M., Gervais, P., Mueller, A., Kossaifi, J., Gramfort, A., Thirion, B., & Varoquaux, G. (2014). Machine learning for neuroimaging with scikit-learn. Frontiers in Neuroinformatics, 8. 10.3389/fninf.2014.00014

Antonenko, D., Külzow, N., Sousa, A., Prehn, K., Grittner, U., & Flöel, A. (2018). Neuronal and behavioral effects of multi-day brain stimulation and memory training. Neurobiology of Aging, 61, 245–254. 10.1016/j.neurobiolaging.2017.09.017

Bakdash, J. Z., & Marusich, L. R. (2017). Repeated Measures Correlation. Frontiers in Psychology, 8. 10.3389/fpsyg.2017.00456

Behzadi, Y., Restom, K., Liau, J., & Liu, T. T. (2007). A component based noise correction method (CompCor) for BOLD and perfusion based fMRI. NeuroImage, 37(1), 90–101. 10.1016/j.neuroimage.2007.04.042

Bellgowan, P. S. F., Buffalo, E. A., Bodurka, J., & Martin, A. (2009). Lateralized spatial and object memory encoding in entorhinal and perirhinal cortices. Learning & Memory, 16(7), 433–438. 10.1101/lm.1357309

Bogdan, P. C., Iordan, A. D., Shobrook, J., & Dolcos, F. (2023). ConnSearch: A framework for functional connectivity analysis designed for interpretability and effectiveness at limited sample sizes. NeuroImage, 278, 120274. 10.1016/j.neuroimage.2023.120274

Buffalo, E. A., Bellgowan, P. S. F., & Martin, A. (2006). Distinct roles for medial temporal lobe structures in memory for objects and their locations. Learning & Memory, 13(5), 638–643. 10.1101/lm.251906

Caviezel, M. P., Reichert, C. F., Sadeghi Bahmani, D., Linnemann, C., Liechti, C., Bieri, O., Borgwardt, S., Leyhe, T., & Melcher, T. (2020). The Neural Mechanisms of Associative Memory Revisited: fMRI Evidence from Implicit Contingency Learning. Frontiers in Psychiatry, 10, 1002. 10.3389/fpsyt.2019.01002

Chai, X. J., Castañón, A. N., Ongür, D., & Whitfield-Gabrieli, S. (2012). Anticorrelations in resting state networks without global signal regression. NeuroImage, 59(2), 1420–1428. 10.1016/j.neuroimage.2011.08.048

Cole, M. W., Bassett, D. S., Power, J. D., Braver, T. S., & Petersen, S. E. (2014). Intrinsic and task-evoked network architectures of the human brain. Neuron, 83(1), 238–251. 10.1016/j.neuron.2014.05.014

Cole, M. W., Ito, T., Cocuzza, C., & Sanchez-Romero, R. (2021). The Functional Relevance of Task-State Functional Connectivity. The Journal of Neuroscience, 41(12), 2684–2702. 10.1523/JNEUROSCI.1713-20.2021

Davachi, L. (2006). Item, context and relational episodic encoding in humans. *Current Opinion in Neurobiology*, Motor Systems / Neurobiology of Behaviour, 16(6), 693–700. 10.1016/j.conb.2006.10.012

De Biase, R., Esposito, S., Chiaramello, E., Parazzini, M., & Sagliano, L. (2025). The role of the hippocampus and retrosplenial cortex in spatial memory: A double blind anodal transcranial direct current stimulation study. Frontiers in Human Neuroscience, 19. 10.3389/fnhum.2025.1661310

de Sousa, A. V. C., Grittner, U., Rujescu, D., Külzow, N., & Flöel, A. (2020). Impact of 3-Day Combined Anodal Transcranial Direct Current Stimulation-Visuospatial Training on Object-Location Memory in Healthy Older Adults and Patients with Mild Cognitive Impairment. Journal of Alzheimer’s Disease: JAD, 75(1), 223–244. 10.3233/JAD-191234

Desikan, R. S., Ségonne, F., Fischl, B., Quinn, B. T., Dickerson, B. C., Blacker, D., Buckner, R. L., Dale, A. M., Maguire, R. P., Hyman, B. T., Albert, M. S., & Killiany, R. J. (2006). An automated labeling system for subdividing the human cerebral cortex on MRI scans into gyral based regions of interest. NeuroImage, 31(3), 968–980. 10.1016/j.neuroimage.2006.01.021

England, H. B., Fyock, C., Meredith Gillis, M., & Hampstead, B. M. (2015). Transcranial direct current stimulation modulates spatial memory in cognitively intact adults. Behavioural Brain Research, 283, 191–195. 10.1016/j.bbr.2015.01.044

Ester-Nacke, T., Berti, K., Veit, R., Dannecker, C., Salvador, R., Ruffini, G., Heni, M., Birkenfeld, A. L., Plewnia, C., Preissl, H., & Kullmann, S. (2024). Network-targeted transcranial direct current stimulation of the hypothalamus appetite-control network: A feasibility study. Scientific Reports, 14(1), 11341. 10.1038/s41598-024-61852-3

Ezzyat, Y., Wanda, P. A., Levy, D. F., Kadel, A., Aka, A., Pedisich, I., Sperling, M. R., Sharan, A. D., Lega, B. C., Burks, A., Gross, R. E., Inman, C. S., Jobst, B. C., Gorenstein, M. A., Davis, K. A., Worrell, G. A., Kucewicz, M. T., Stein, J. M., Gorniak, R., … Kahana, M. J. (2018). Closed-loop stimulation of temporal cortex rescues functional networks and improves memory. Nature Communications, 9, 365. 10.1038/s41467-017-02753-0

Fair, D. A., Schlaggar, B. L., Cohen B.A., A. L., Miezin, F. M., Dosenbach, N. U. F., Wenger, K. K., Fox, M. D., Snyder, A. Z., Raichle, M. E., & Petersen, S. E. (2007). A METHOD FOR USING BLOCKED AND EVENT-RELATED FMRI DATA TO STUDY “RESTING STATE” FUNCTIONAL CONNECTIVITY. NeuroImage, 35(1), 396–405. 10.1016/j.neuroimage.2006.11.051

Fiori, V., Kunz, L., Kuhnke, P., Marangolo, P., & Hartwigsen, G. (2018). Transcranial direct current stimulation (tDCS) facilitates verb learning by altering effective connectivity in the healthy brain. NeuroImage, 181, 550–559. 10.1016/j.neuroimage.2018.07.040

Flöel, A., Suttorp, W., Kohl, O., Kürten, J., Lohmann, H., Breitenstein, C., & Knecht, S. (2012). Non-invasive brain stimulation improves object-location learning in the elderly. Neurobiology of Aging, 33(8), Article 8. 10.1016/j.neurobiolaging.2011.05.007

Fox, M. D., Buckner, R. L., Liu, H., Chakravarty, M. M., Lozano, A. M., & Pascual-Leone, A. (2014). Resting-state networks link invasive and noninvasive brain stimulation across diverse psychiatric and neurological diseases. Proceedings of the National Academy of Sciences of the United States of America, 111(41), E4367–4375. 10.1073/pnas.1405003111

Frazier, J. A., Chiu, S., Breeze, J. L., Makris, N., Lange, N., Kennedy, D. N., Herbert, M. R., Bent, E. K., Koneru, V. K., Dieterich, M. E., Hodge, S. M., Rauch, S. L., Grant, P. E., Cohen, B. M., Seidman, L. J., Caviness, V. S., & Biederman, J. (2005). Structural brain magnetic resonance imaging of limbic and thalamic volumes in pediatric bipolar disorder. The American Journal of Psychiatry, 162(7), 1256–1265. 10.1176/appi.ajp.162.7.1256

Friston, K. J., Buechel, C., Fink, G. R., Morris, J., Rolls, E., & Dolan, R. J. (1997). Psychophysiological and modulatory interactions in neuroimaging. NeuroImage, 6(3), 218–229. 10.1006/nimg.1997.0291

Friston, K. J., Williams, S., Howard, R., Frackowiak, R. S., & Turner, R. (1996). Movement-related effects in fMRI time-series. Magnetic Resonance in Medicine, 35(3), 346–355. 10.1002/mrm.1910350312

Fromm, A. E., Grittner, U., Brodt, S., Flöel, A., & Antonenko, D. (2024). No Object–Location Memory Improvement through Focal Transcranial Direct Current Stimulation over the Right Temporoparietal Cortex. Life, 14(5), Article 5. 10.3390/life14050539

Gillis, M. M., Garcia, S., & Hampstead, B. M. (2016). Working memory contributes to the encoding of object location associations: Support for a 3-part model of object location memory. Behavioural Brain Research, 311, 192–200. 10.1016/j.bbr.2016.05.037

Goicoechea, E. B., Agres, P. F., Rau, J. M., Agustín, A. S., & Voss, J. L. (2025). A meta-analysis suggests that TMS targeting the hippocampal network selectively improves episodic memory. eLife, 14. 10.7554/eLife.108934.1

Goldstein, J. M., Seidman, L. J., Makris, N., Ahern, T., O’Brien, L. M., Caviness, V. S., Kennedy, D. N., Faraone, S. V., & Tsuang, M. T. (2007). Hypothalamic abnormalities in schizophrenia: Sex effects and genetic vulnerability. Biological Psychiatry, 61(8), 935–945. 10.1016/j.biopsych.2006.06.027

Grill-Spector, K., Sayres, R., & Ress, D. (2006). High-resolution imaging reveals highly selective nonface clusters in the fusiform face area. Nature Neuroscience, 9(9), 1177–1185. 10.1038/nn1745

Hales, J. B., & Brewer, J. B. (2013). Parietal and frontal contributions to episodic encoding of location. Behavioural Brain Research, 243, 16–20. 10.1016/j.bbr.2012.12.048

Hallquist, M. N., Hwang, K., & Luna, B. (2013). The nuisance of nuisance regression: Spectral misspecification in a common approach to resting-state fMRI preprocessing reintroduces noise and obscures functional connectivity. NeuroImage, 82, 208–225. 10.1016/j.neuroimage.2013.05.116

Hannula, D. E., & Ranganath, C. (2008). Medial Temporal Lobe Activity Predicts Successful Relational Memory Binding. The Journal of Neuroscience, 28(1), 116–124. 10.1523/JNEUROSCI.3086-07.2008

Hayes, S. M., Ryan, L., Schnyer, D. M., & Nadel, L. (2004). An fMRI Study of Episodic Memory: Retrieval of Object, Spatial, and Temporal Information. Behavioral Neuroscience, 118(5), 885–896. 10.1037/0735-7044.118.5.885

Hedden, T., & Gabrieli, J. D. E. (2004). Insights into the ageing mind: A view from cognitive neuroscience. Nature Reviews. Neuroscience, 5(2), 87–96. 10.1038/nrn1323

Jeong, W., Chung, C. K., & Kim, J. S. (2015). Episodic memory in aspects of large-scale brain networks. Frontiers in Human Neuroscience, 9. 10.3389/fnhum.2015.00454

Karolis, V. R., Corbetta, M., & Thiebaut de Schotten, M. (2019). The architecture of functional lateralisation and its relationship to callosal connectivity in the human brain. Nature Communications, 10(1), 1417. 10.1038/s41467-019-09344-1

Kessels, R. P. C., Hobbel, D., & Postma, A. (2007). Aging, context memory and binding: A comparison of “what, where and when” in young and older adults. International Journal of Neuroscience, 117(6), 795–810. 10.1080/00207450600910218

Kessels, R. P. C., Rijken, S., Joosten-Weyn Banningh, L. W. A., Van Schuylenborgh-VAN Es, N., & Olde Rikkert, M. G. M. (2010). Categorical spatial memory in patients with mild cognitive impairment and Alzheimer dementia: Positional versus object-location recall. Journal of the International Neuropsychological Society: JINS, 16(1), 200–204. 10.1017/S1355617709990944

Kocataş, H., Abdelmotaleb, M., Guevara, L. M. C., Niemann, F., Shahbabaie, A., Malinowski, R., Riemann, S., Hayek, D., Antonenko, D., Rodriguez-Fornells, A., Flöel, A., & Meinzer, M. (2025). Functionally Relevant and Reliable Brain Stimulation Targets for Enhancement of Novel Word-Learning (p. 2025.11.04.686295). bioRxiv. 10.1101/2025.11.04.686295

Kuhnke, P., Kiefer, M., & Hartwigsen, G. (2021). Task-Dependent Functional and Effective Connectivity during Conceptual Processing. Cerebral Cortex, 31(7), 3475–3493. 10.1093/cercor/bhab026

Kuhnke, P., Numssen, O., Voeller, J., Cheung, V. K. M., Weise, K., Kiefer, M., & Hartwigsen, G. (2025). Left inferior parietal lobe and auditory cortex jointly contribute to sound knowledge retrieval. Brain Stimulation, 18(4), 1037–1047. 10.1016/j.brs.2025.05.113

Külzow, N., Kerti, L., Witte, V. A., Kopp, U., Breitenstein, C., & Flöel, A. (2014). An object location memory paradigm for older adults with and without mild cognitive impairment. Journal of Neuroscience Methods, 237, 16–25. 10.1016/j.jneumeth.2014.08.020

Lee, A. C. H., Rahman, S., Hodges, J. R., Sahakian, B. J., & Graham, K. S. (2003). Associative and recognition memory for novel objects in dementia: Implications for diagnosis. The European Journal of Neuroscience, 18(6), 1660–1670. 10.1046/j.1460-9568.2003.02883.x

Liu, X., Qiu, F., Hou, L., & Wang, X. (2022). Review of Noninvasive or Minimally Invasive Deep Brain Stimulation. Frontiers in Behavioral Neuroscience, 15. 10.3389/fnbeh.2021.820017

Lynch, L. K., Lu, K., Wen, H., Zhang, Y., Saykin, A. J., & Liu, Z. (2018). Task-evoked functional connectivity does not explain functional connectivity differences between rest and task conditions. Human Brain Mapping, 39(12), 4939–4948. 10.1002/hbm.24335

Makris, N., Goldstein, J. M., Kennedy, D., Hodge, S. M., Caviness, V. S., Faraone, S. V., Tsuang, M. T., & Seidman, L. J. (2006). Decreased volume of left and total anterior insular lobule in schizophrenia. Schizophrenia Research, 83(2–3), 155–171. 10.1016/j.schres.2005.11.020

Masharipov, R., Knyazeva, I., Korotkov, A., Cherednichenko, D., & Kireev, M. (2024). Comparison of whole-brain task-modulated functional connectivity methods for fMRI task connectomics. Communications Biology, 7(1), 1402. 10.1038/s42003-024-07088-3

McLaren, D. G., Ries, M. L., Xu, G., & Johnson, S. C. (2012). A Generalized Form of Context-Dependent Psychophysiological Interactions (gPPI): A Comparison to Standard Approaches. Neuroimage, 61(4), 1277–1286. 10.1016/j.neuroimage.2012.03.068

Meinzer, M., Antonenko, D., Lindenberg, R., Hetzer, S., Ulm, L., Avirame, K., Flaisch, T., & Flöel, A. (2012). Electrical Brain Stimulation Improves Cognitive Performance by Modulating Functional Connectivity and Task-Specific Activation. Journal of Neuroscience, 32(5), 1859–1866. 10.1523/JNEUROSCI.4812-11.2012

Meinzer, M., Shahbabaie, A., Antonenko, D., Blankenburg, F., Fischer, R., Hartwigsen, G., Nitsche, M. A., Li, S.-C., Thielscher, A., Timmann, D., Waltemath, D., Abdelmotaleb, M., Kocataş, H., Caisachana Guevara, L. M., Batsikadze, G., Grundei, M., Cunha, T., Hayek, D., Turker, S., … Flöel, A. (2024). Investigating the neural mechanisms of transcranial direct current stimulation effects on human cognition: Current issues and potential solutions. Frontiers in Neuroscience, 18, 1389651. 10.3389/fnins.2024.1389651

Montaldi, D., & Mayes, A. R. (2010). The role of recollection and familiarity in the functional differentiation of the medial temporal lobes. Hippocampus, 20(11), 1291–1314. 10.1002/hipo.20853

Morfini, F., Whitfield-Gabrieli, S., & Nieto-Castañón, A. (2023). Functional connectivity MRI quality control procedures in CONN. Frontiers in Neuroscience, 17. 10.3389/fnins.2023.1092125

Niemann, F., Riemann, S., Hubert, A.-K., Antonenko, D., Thielscher, A., Martin, A. K., Unger, N., Flöel, A., & Meinzer, M. (2024). Electrode positioning errors reduce current dose for focal tDCS set-ups: Evidence from individualized electric field mapping. Clinical Neurophysiology, 162, 201–209. 10.1016/j.clinph.2024.03.031

Nieto-Castanon, A. (2020). Handbook of functional connectivity Magnetic Resonance Imaging methods in CONN. 10.56441/hilbertpress.2207.6598

Nieto-Castanon, A., & Whitfield-Gabrieli, S. (2022). CONN functional connectivity toolbox: RRID SCR_009550, release 22. 10.56441/hilbertpress.2246.5840

Nilakantan, A. S., Bridge, D. J., Gagnon, E. P., VanHaerents, S. A., & Voss, J. L. (2017). Stimulation of the posterior cortical-hippocampal network enhances precision of memory recollection. Current Biology : CB, 27(3), 465–470. 10.1016/j.cub.2016.12.042

Nilearn contributors. (n.d.). Nilearn [Computer software]. Retrieved https://github.com/nilearn/nilearn

Passiatore, R., Lupo, A., Sambuco, N., Antonucci, L. A., Stolfa, G., Bertolino, A., Popolizio, T., Suchan, B., & Pergola, G. (2025). Interindividual Variability in Memory Performance Is Related to Corticothalamic Networks during Memory Encoding and Retrieval. The Journal of Neuroscience, 45(19), e0975242025. 10.1523/JNEUROSCI.0975-24.2025

Penny, W., Friston, K., Ashburner, J., Kiebel, S., & Nichols, T. (2007). Statistical Parametric Mapping: The Analysis of Functional Brain Images. 10.1016/B978-0-12-372560-8.X5000-1

Persson, J., Stening, E., Nordin, K., & Söderlund, H. (2018). Predicting episodic and spatial memory performance from hippocampal resting-state functional connectivity: Evidence for an anterior-posterior division of function. Hippocampus, 28(1), 53–66. 10.1002/hipo.22807

Petersson, K. M., Sandblom, J., Gisselgård, J., & Ingvar, M. (2001). Learning related modulation of functional retrieval networks in man. Scandinavian Journal of Psychology, 42(3), 197–216. 10.1111/1467-9450.00231

Postma, A., Kessels, R. P. C., & van Asselen, M. (2004). The Neuropsychology of Object-Location Memory. In Human spatial memory: Remembering where. (pp. 143–160). Lawrence Erlbaum Associates Publishers.

Postma, A., Kessels, R. P. C., & van Asselen, M. (2008). How the brain remembers and forgets where things are: The neurocognition of object-location memory. Neuroscience and Biobehavioral Reviews, 32(8), 1339–1345. 10.1016/j.neubiorev.2008.05.001

Power, J. D., Barnes, K. A., Snyder, A. Z., Schlaggar, B. L., & Petersen, S. E. (2012). Spurious but systematic correlations in functional connectivity MRI networks arise from subject motion. NeuroImage, 59(3), 2142–2154. 10.1016/j.neuroimage.2011.10.018

Power, J. D., Mitra, A., Laumann, T. O., Snyder, A. Z., Schlaggar, B. L., & Petersen, S. E. (2014). Methods to detect, characterize, and remove motion artifact in resting state fMRI. NeuroImage, 84, 320–341. 10.1016/j.neuroimage.2013.08.048

Prehn, K., Stengl, H., Grittner, U., Kosiolek, R., Ölschläger, A., Weidemann, A., & Flöel, A. (2017). Effects of Anodal Transcranial Direct Current Stimulation and Serotonergic Enhancement on Memory Performance in Young and Older Adults. Neuropsychopharmacology: Official Publication of the American College of Neuropsychopharmacology, 42(2), 551–561. 10.1038/npp.2016.170

Ritchey, M., Yonelinas, A. P., & Ranganath, C. (2014). Functional connectivity relationships predict similarities in task activation and pattern information during associative memory encoding. Journal of Cognitive Neuroscience, 26(5), 1085–1099. 10.1162/jocn_a_00533

Rolls, E. T. (2021). Learning Invariant Object and Spatial View Representations in the Brain Using Slow Unsupervised Learning. Frontiers in Computational Neuroscience, 15, 686239. 10.3389/fncom.2021.686239

Rolls, E. T. (2023). Hippocampal spatial view cells for memory and navigation, and their underlying connectivity in humans. Hippocampus, 33(5), 533–572. 10.1002/hipo.23467

Rolls, E. T. (2024). Two what, two where, visual cortical streams in humans. Neuroscience and Biobehavioral Reviews, 160, 105650. 10.1016/j.neubiorev.2024.105650

Rolls, Edmund T, Deco, G., Huang, C.-C., & Feng, J. (2023). Multiple cortical visual streams in humans. Cerebral Cortex, 33(7), 3319–3349. 10.1093/cercor/bhac276

Rolls, Edmund T., Deco, G., Zhang, Y., & Feng, J. (2023). Hierarchical organization of the human ventral visual streams revealed with magnetoencephalography. Cerebral Cortex (New York, N.Y.: 1991), 33(20), 10686–10701. 10.1093/cercor/bhad318

Rolls, E. T., & Treves, A. (2024). A theory of hippocampal function: New developments. Progress in Neurobiology, 238, 102636. 10.1016/j.pneurobio.2024.102636

Rolls, E. T., Yan, X., Deco, G., Zhang, Y., Jousmaki, V., & Feng, J. (2024). A ventromedial visual cortical ‘Where’ stream to the human hippocampus for spatial scenes revealed with magnetoencephalography. Communications Biology, 7(1), 1047. 10.1038/s42003-024-06719-z

Rolls, E. T., Zhang, R., Deco, G., Vatansever, D., & Feng, J. (2024). Selective Brain Activations and Connectivities Related to the Storage and Recall of Human Object-Location, Reward-Location, and Word-Pair Episodic Memories. Human Brain Mapping, 45(15), e70056. 10.1002/hbm.70056

Sadaghiani, S., & D’Esposito, M. (2015). Functional Characterization of the Cingulo-Opercular Network in the Maintenance of Tonic Alertness. *Cerebral Cortex (New York*, NY), 25(9), 2763–2773. 10.1093/cercor/bhu072

Satake, T., Taki, A., Kasahara, K., Yoshimaru, D., & Tsurugizawa, T. (2024). Comparison of local activation, functional connectivity, and structural connectivity in the N-back task. Frontiers in Neuroscience, 18. 10.3389/fnins.2024.1337976

Schedlbauer, A. M., Copara, M. S., Watrous, A. J., & Ekstrom, A. D. (2014). Multiple interacting brain areas underlie successful spatiotemporal memory retrieval in humans. Scientific Reports, 4(1), 6431. 10.1038/srep06431

Seabold, S., & Perktold, J. (2010). Statsmodels: Econometric and Statistical Modeling with Python. SciPy 2010. 10.25080/Majora-92bf1922-011

Shahbabaie, A., Abdelmotaleb, M., Kocataş, H., Niemann, F., Antonenko, D., Flöel, A., & Meinzer, M. (2026). Multimodal Imaging-Based Targeting Approach for Network-Level Brain Stimulation (p. 2026.01.09.698588). bioRxiv. 10.64898/2026.01.09.698588

Shi, Y., Yang, L., Lu, J., Yan, T., Ding, Y., & Wang, B. (2024). The dynamic reconfiguration of the functional network during episodic memory task predicts the memory performance. Scientific Reports, 14(1), 20527. 10.1038/s41598-024-71295-5

Siebner, H. R., Funke, K., Aberra, A. S., Antal, A., Bestmann, S., Chen, R., Classen, J., Davare, M., Di Lazzaro, V., Fox, P. T., Hallett, M., Karabanov, A. N., Kesselheim, J., Beck, M. M., Koch, G., Liebetanz, D., Meunier, S., Miniussi, C., Paulus, W., … Ugawa, Y. (2022). Transcranial magnetic stimulation of the brain: What is stimulated? – A consensus and critical position paper. Clinical Neurophysiology : Official Journal of the International Federation of Clinical Neurophysiology, 140, 59–97. 10.1016/j.clinph.2022.04.022

Sliwinska, M. W., Violante, I. R., Wise, R. J. S., Leech, R., Devlin, J. T., Geranmayeh, F., & Hampshire, A. (2017). Stimulating Multiple-Demand Cortex Enhances Vocabulary Learning. Journal of Neuroscience, 37(32), 7606–7618. 10.1523/JNEUROSCI.3857-16.2017

Squire, L. R., & Wixted, J. T. (2011). The Cognitive Neuroscience of Human Memory Since H.M. Annual Review of Neuroscience, 34, 259–288. 10.1146/annurev-neuro-061010-113720

Staresina, B. P., Michelmann, S., Bonnefond, M., Jensen, O., Axmacher, N., & Fell, J. (2016). Hippocampal pattern completion is linked to gamma power increases and alpha power decreases during recollection. eLife, 5, e17397. 10.7554/eLife.17397

Sučević, J., & Schapiro, A. C. (2023). A neural network model of hippocampal contributions to category learning. eLife, 12, e77185. 10.7554/eLife.77185

Tambini, A., Ketz, N., & Davachi, L. (2010). Enhanced brain correlations during rest are related to memory for recent experiences. Neuron, 65(2), 280–290. 10.1016/j.neuron.2010.01.001

Tang, Y., Xu, L., Zhu, T., Cui, H., Qian, Z., Kong, G., Tang, X., Wei, Y., Zhang, T., Hu, Y., Sheng, J., & Wang, J. (2023). Visuospatial Learning Selectively Enhanced by Personalized Transcranial Magnetic Stimulation over Parieto-Hippocampal Network among Patients at Clinical High-Risk for Psychosis. Schizophrenia Bulletin, 49(4), 923–932. 10.1093/schbul/sbad015

Thair, H., Holloway, A. L., Newport, R., & Smith, A. D. (2017). Transcranial Direct Current Stimulation (tDCS): A Beginner’s Guide for Design and Implementation. Frontiers in Neuroscience, 11, 641. 10.3389/fnins.2017.00641

Thielscher, A., Hayek, D., Puonti, O., Grittner, U., Blankenburg, F., Fischer, R., Hartwigsen, G., Li, S.-C., Meinzer, M., Nitsche, M. A., Timmann, D., Flöel, A., & Antonenko, D. (2026). Harmonizing the stimulation dose of focal tDCS across target sites (p. 2026.01.09.698549). bioRxiv. 10.64898/2026.01.09.698549

Tuominen, J., Specht, K., Vaisvilaite, L., & Zeidman, P. (2023). An information-theoretic analysis of resting-state versus task fMRI. Network Neuroscience, 7(2), 769–786. 10.1162/netn_a_00302

Ungerleider, L. G., & Haxby, J. V. (1994). “What” and “where” in the human brain. Current Opinion in Neurobiology, 4(2), 157–165. 10.1016/0959-4388(94)90066-3

Vallat, R. (2018). Pingouin: Statistics in Python. Journal of Open Source Software, 3(31), 1026. 10.21105/joss.01026

Virtanen, P., Gommers, R., Oliphant, T. E., Haberland, M., Reddy, T., Cournapeau, D., Burovski, E., Peterson, P., Weckesser, W., Bright, J., van der Walt, S. J., Brett, M., Wilson, J., Millman, K. J., Mayorov, N., Nelson, A. R. J., Jones, E., Kern, R., Larson, E., … van Mulbregt, P. (2020). SciPy 1.0: Fundamental algorithms for scientific computing in Python. Nature Methods, 17(3), 261–272. 10.1038/s41592-019-0686-2

Vossel, S., Geng, J. J., & Fink, G. R. (2014). Dorsal and Ventral Attention Systems. The Neuroscientist, 20(2), 150–159. 10.1177/1073858413494269

Wander, J. D., Blakely, T., Miller, K. J., Weaver, K. E., Johnson, L. A., Olson, J. D., Fetz, E. E., Rao, R. P. N., & Ojemann, J. G. (2013). Distributed cortical adaptation during learning of a brain–computer interface task. Proceedings of the National Academy of Sciences, 110(26), 10818–10823. 10.1073/pnas.1221127110

Wang, J. X., Rogers, L. M., Gross, E. Z., Ryals, A. J., Dokucu, M. E., Brandstatt, K. L., Hermiller, M. S., & Voss, J. L. (2014). Targeted enhancement of cortical-hippocampal brain networks and associative memory. Science, 345(6200), 1054–1057. 10.1126/science.1252900

Warren, K. N., Hermiller, M. S., Nilakantan, A. S., & Voss, J. L. (2019). Stimulating the hippocampal posterior-medial network enhances task-dependent connectivity and memory. eLife, 8, e49458. 10.7554/eLife.49458

Whitfield-Gabrieli, S., & Nieto-Castanon, A. (2012). Conn: A functional connectivity toolbox for correlated and anticorrelated brain networks. Brain Connectivity, 2(3), 125–141. 10.1089/brain.2012.0073

Whitfield-Gabrieli, S., Nieto-Castanon, A., & Ghosh, S. (2011). Artifact detection tools (ART). *Cambridge*, MA. Release Version, 7(19), 11.

