## Supplementary for "Temporo-Occipital and Medial Temporal Networks Underlying Object-Location Learning"

### Supplementary Materials

**Quality Control (QC)**

Head motion and other artifacts can markedly bias fMRI connectivity metrics (Power et al., 2012). To mitigate these effects, we implemented a rigorous, stepwise QC procedure that covered raw-data screening, verification of preprocessing outputs, and assessment of denoised time series, following the recommended pipeline (Nieto-Castanon & Whitfield-Gabrieli, 2022; Morfini et al., 2023).

For the preprocessing step, all functional and anatomical images underwent visual inspection to assess spatial normalization, segmentation accuracy, and anatomical-functional alignment. Automated QC metrics were also extracted from preprocessing outputs. These included measures of motion (maximum and mean framewise displacement; FWD), global signal change (GSC), and the number and proportion of valid versus outlier scans per run. Additional normalization quality metrics (NORMfunc and NORManat) quantified the similarity between each participant's normalized gray matter mask and the MNI template.

For denoising, both visual inspection and quantitative quality control (QC) metrics were employed to ensure the integrity of the BOLD signal. Each participant's BOLD time series was reviewed to identify potential signs of over- or under-correction, including inspection of carpet plots (Power, 2017) before and after denoising, alongside traces of framewise displacement (FWD), global signal change (GSC), and time points flagged as outliers. Quantitative QC measures were then used to evaluate post-denoising data quality, including the effective degrees of freedom (DOF), which reflects the remaining usable signal; BOLD standard deviation (BOLDstd), indicating signal stability; and global correlation (GCOR), which captures the average correlation across all brain voxels to identify residual global noise (Saad et al., 2013). Outliers were defined as values exceeding Q3 (3$^{rd}$quartile) + 3×IQR (interquartile range) or below Q1 (1$^{st}$ quartile) − 3×IQR.

Functional connectivity (FC) was estimated via pairwise correlations among 1,000 randomly sampled gray matter voxels (Morfini et al., 2023). One participant with FC distributions deviating from the group norm was flagged. To quantify the influence of motion and scanner-related artifacts on FC, the QC-FC% metric was calculated (Ciric et al., 2017), indicating the proportion of functional connections significantly associated with head motion or other data quality indices. A strict quality criterion required QC-FC% ≥ 95%. Participants identified as outliers on one or more QC metrics or FC distributions were considered for exclusion using a stepwise approach to maximize sample retention while meeting the QC-FC% threshold. Excluding one participant was sufficient to achieve the 95% criterion post-denoising, resulting in 19 participants in the final analysis (see below Supplementary Figure S1).

**Figure S1:**

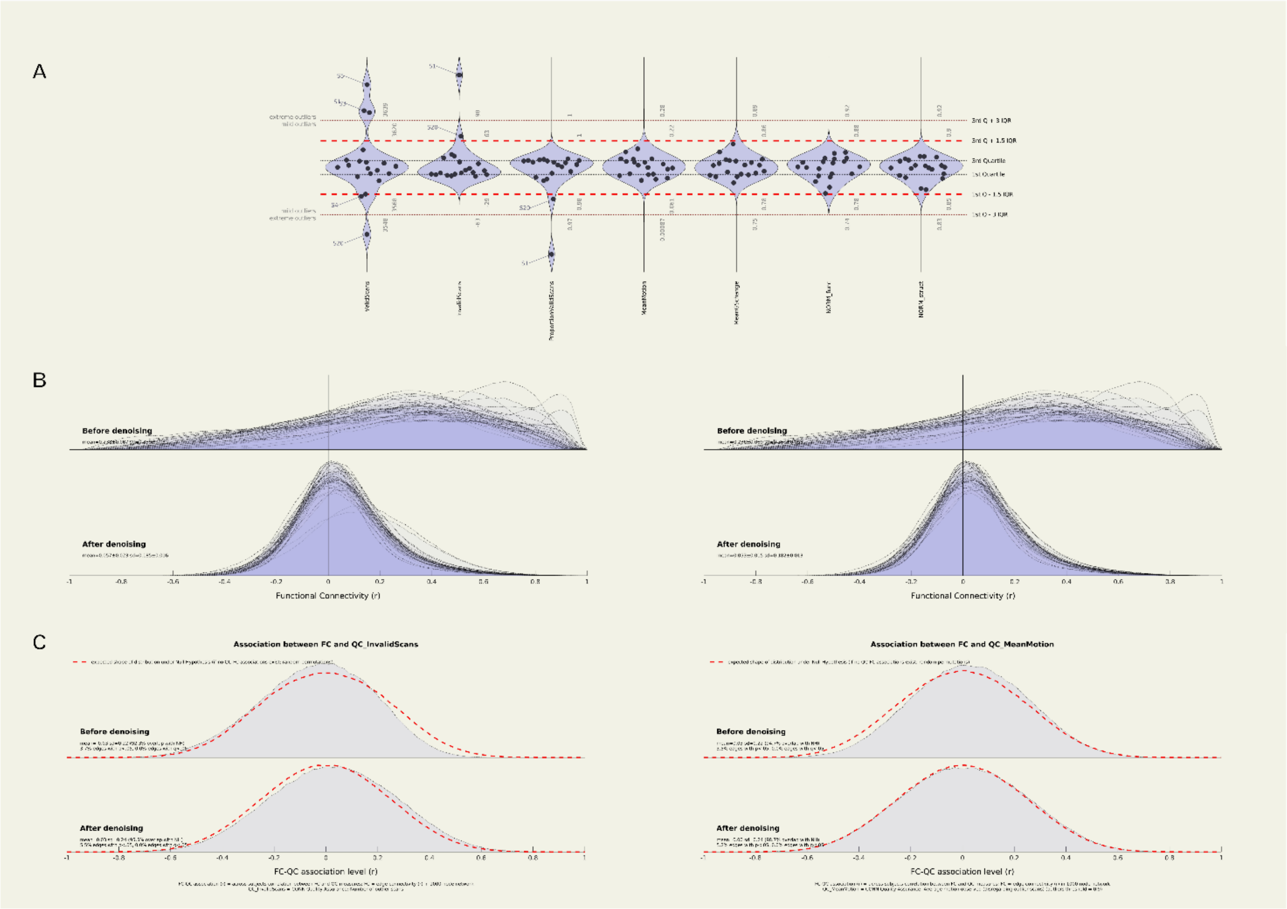

**Figure S1. Quality control (QC) metrics for functional connectivity analysis.**
(A) Individual QC measures for all 20 participants prior to exclusion, including valid scans, invalid scans, mean motion, mean global signal change, and normalization quality for structural and functional images. Boundary lines indicate thresholds for mild and extreme outliers. (B) Distributions of functional connectivity (FC) estimates before (upper) and after (lower) denoising. Plots are shown for all 20 participants (left) and following exclusion of one participant who deviated from the group norm (right). (C) Associations between FC strength and data quality indices (QC-FC%) before and after denoising. The left panel shows QC-FC correlations with invalid scans, and the right panel shows correlations with mean motion. Upper curves represent values before denoising, and lower curves represent values after denoising.

**Seed-to-voxel analyses**

Additional seed-to-voxel analyses were subsequently performed to complement and validate the ROI-to-ROI findings and to characterize whole-brain functional connectivity patterns associated with each OLM-relevant seed region.

Seed regions were defined using two complementary strategies. First, anatomically defined ROIs were selected a priori based on the regions consistently implicated in OLM. Second, functionally defined seed regions were created by placing 5-mm-radius spherical ROIs at the peak MNI coordinates of activation clusters that showed significant differences between the Learning and Control conditions in the univariate activation analysis (Abdelmotaleb et al., 2025). Both seed-definition approaches yielded highly similar connectivity patterns.

Across anatomical seeds, OLM learning was associated with a robust and coherent connectivity pattern linking MTL structures with ventral visual and occipito-temporal cortices. Significant connectivity maps are presented below in Figure S2. Specifically, hippocampal and parahippocampal seeds exhibited increased coupling with bilateral inferior and middle temporal gyri (temporo-occipital portions), fusiform gyri, and lateral occipital cortices, indicating enhanced integration between memory-related regions and high-level visual processing areas during learning. This pattern was largely bilateral and showed substantial interhemispheric homologue connectivity. Seeds placed in occipito-temporal regions (middle and inferior temporal gyri and lateral occipital cortices) similarly demonstrated strengthened reciprocal connectivity within the ventral visual stream, extending to MTL regions, including the hippocampus and parahippocampus.

In contrast, connectivity patterns observed during the control condition differed markedly. Across several seeds, most prominently the parahippocampal and lateral occipital cortices, stronger connectivity during control was observed with regions associated with domain-general attentional, salience, and sensorimotor processing, including the cingulate and paracingulate cortices, precuneus, frontal poles, insula, operculum, cuneus, and sensorimotor areas.

**Figure S2**

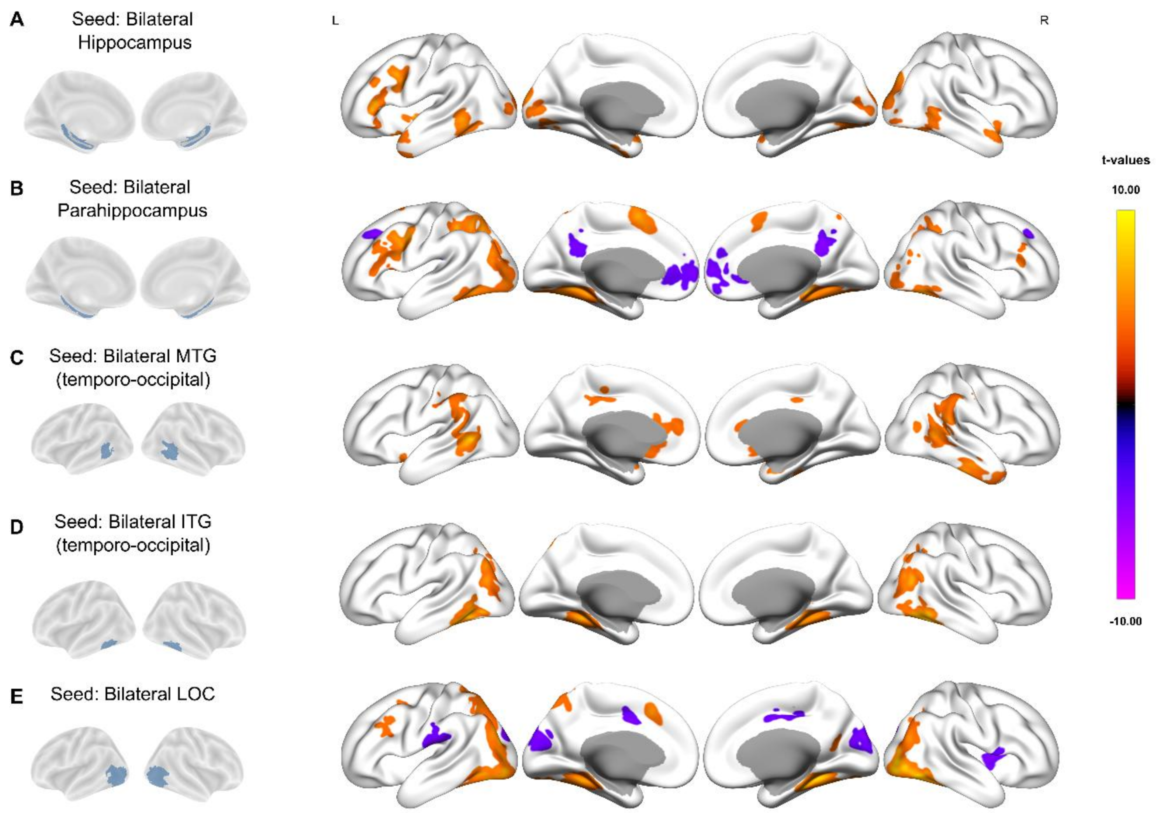

**Figure S2. Surface-rendered group statistical maps showing seed-to-voxel gPPI connectivity for the learning > control contrast of the object-location memory (OLM) task across participants (n = 19).** Maps are displayed on an inflated cortical surface template for the following Harvard-Oxford atlas seeds (top to bottom rows): A. bilateral hippocampi, B. bilateral parahippocampal gyri, C. bilateral middle temporal gyri (MTG, temporo-occipital part), D. bilateral inferior temporal gyri (ITG, temporo-occipital part), E. bilateral lateral occipital cortices (LOC). Each row shows lateral and medial views of both hemispheres (L = left, R = right). Statistical maps were thresholded at $p < 0.001$ with cluster-level family-wise error correction (FWEc) at $p < 0.05$, and bilateral seeds results were shown for illustration purposes. Color bars to the right of each row indicate the t-value scale for that seed: warm colors represent positive t-values (stronger connectivity during learning than control), whereas cool colors represent negative t-values (stronger connectivity during control than learning). Significant clusters and peak MNI coordinates for each seed are reported in supplementary tables (S1-S10).

**Activation ROI-based seed results:**

We report here the functional connectivity results of 5-mm radius sphere ROIs derived from the peak MNI coordinates of significant clusters identified in the univariate activation analysis. Full statistical details are provided in Supplementary Tables S11-S16 and Figure S3.

*Bilateral Temporal-occipital fusiform cortex*

The largest cluster in the univariate activation analysis was located in the right temporal-occipital fusiform gyrus. The sphere ROI centered on the peak MNI coordinate of this cluster showed greater functional connectivity during OLM learning with the bilateral parahippocampal gyri, bilateral temporo-occipital portions of the inferior temporal gyri, bilateral lateral occipital cortices, the right thalamus, and the cerebellum. In contrast, during the control task, the right temporal-occipital fusiform gyrus was primarily connected with the right precentral gyrus, right postcentral gyrus, and right central opercular cortex.

Similarly, the left temporal-occipital fusiform gyrus exhibited stronger connectivity during learning with the bilateral parahippocampal gyri, bilateral occipital and posterior divisions of the fusiform gyri, bilateral lateral occipital cortices, bilateral temporo-occipital inferior temporal gyri, the left paracingulate gyrus, and the cerebellum. During the control task, greater connectivity was observed with the bilateral lingual gyri, right cuneal cortex, and right intracalcarine and supracalcarine cortices.

*Bilateral Temporal fusiform, posterior division cortex*

The posterior division of the right temporal fusiform cortex exhibited greater connectivity during learning with the bilateral fusiform gyri, right parahippocampal gyrus, bilateral lateral occipital cortices, bilateral lingual gyri, bilateral temporo-occipital inferior temporal gyri, the left middle and inferior frontal gyri, and the cerebellum. During the control task, stronger connectivity was observed with the cingulate gyrus, left angular gyrus, left precentral gyrus, and left lateral occipital cortex.

The posterior division of the left temporal fusiform cortex showed stronger connectivity during learning with the bilateral parahippocampal gyri, bilateral fusiform gyri, bilateral lateral occipital cortices, left precentral gyrus, bilateral lingual gyri, bilateral superior parietal lobules, the left superior, middle, and inferior frontal gyri, and the cerebellum. During the control task, it was more strongly connected with the anterior and posterior divisions of the cingulate gyrus, bilateral paracingulate gyri, bilateral frontal poles, and the precuneus cortex.

*Bilateral Lateral occipital cortices (LOCs)*

The right lateral occipital cortex was more functionally connected during the learning task with its contralateral homologue, bilateral parahippocampal gyri, right temporo-occipital inferior temporal gyrus, precuneus cortex, bilateral fusiform gyri, bilateral lingual gyri, and the cerebellum. During the control task, stronger connectivity was observed with the bilateral supramarginal gyri, bilateral postcentral gyri, left parietal operculum, bilateral occipital poles, left middle temporal gyrus, and right frontal pole. The left lateral occipital cortex was more strongly connected during learning with the bilateral hippocampal gyri, fusiform gyri, contralateral lateral occipital cortex, precuneus cortex, and the right temporo-occipital inferior temporal gyrus. During the control task, it showed greater connectivity with the right supramarginal gyrus, right frontal pole, and right insula.

**Figure S3**

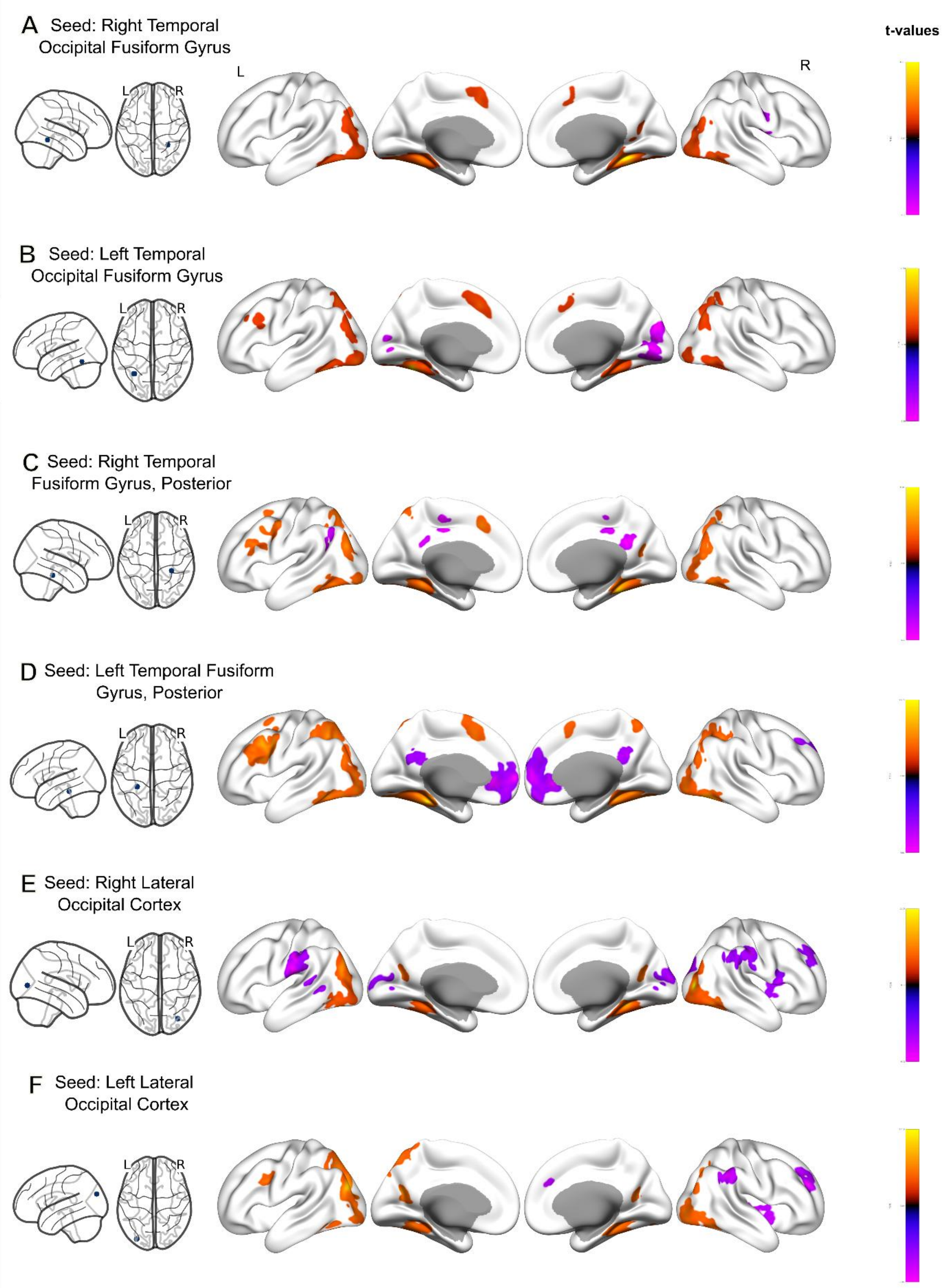

**Figure S3. Surface-rendered group statistical maps showing seed-to-voxel gPPI connectivity for the Learning > Control contrast of the object-location memory (OLM) task across participants (n = 19).**

Maps are displayed on an inflated cortical surface template for the following functionally defined seeds (top to bottom rows): right (A) and left (B) temporal-occipital fusiform gyrus, right (C) and left (D) temporal fusiform gyrus (posterior division), and right (E) and left (F) lateral occipital cortex. Seeds were defined by placing 5-mm radius spheres at the peak MNI coordinates of activation clusters that showed significant differences between Learning and Control conditions in the univariate activation analysis. Each row shows lateral and medial views of both hemispheres (L = left, R = right). Statistical maps were thresholded at $p < 0.001$, with cluster-level family-wise error correction (FWEc). Color bars to the right of each row indicate the t-value scale for that seed: warm colors represent stronger connectivity during Learning than Control, whereas cool colors represent the opposite pattern. Significant clusters and peak MNI coordinates for each seed are reported in Supplementary Tables S11-S16.

**Figure S4:**

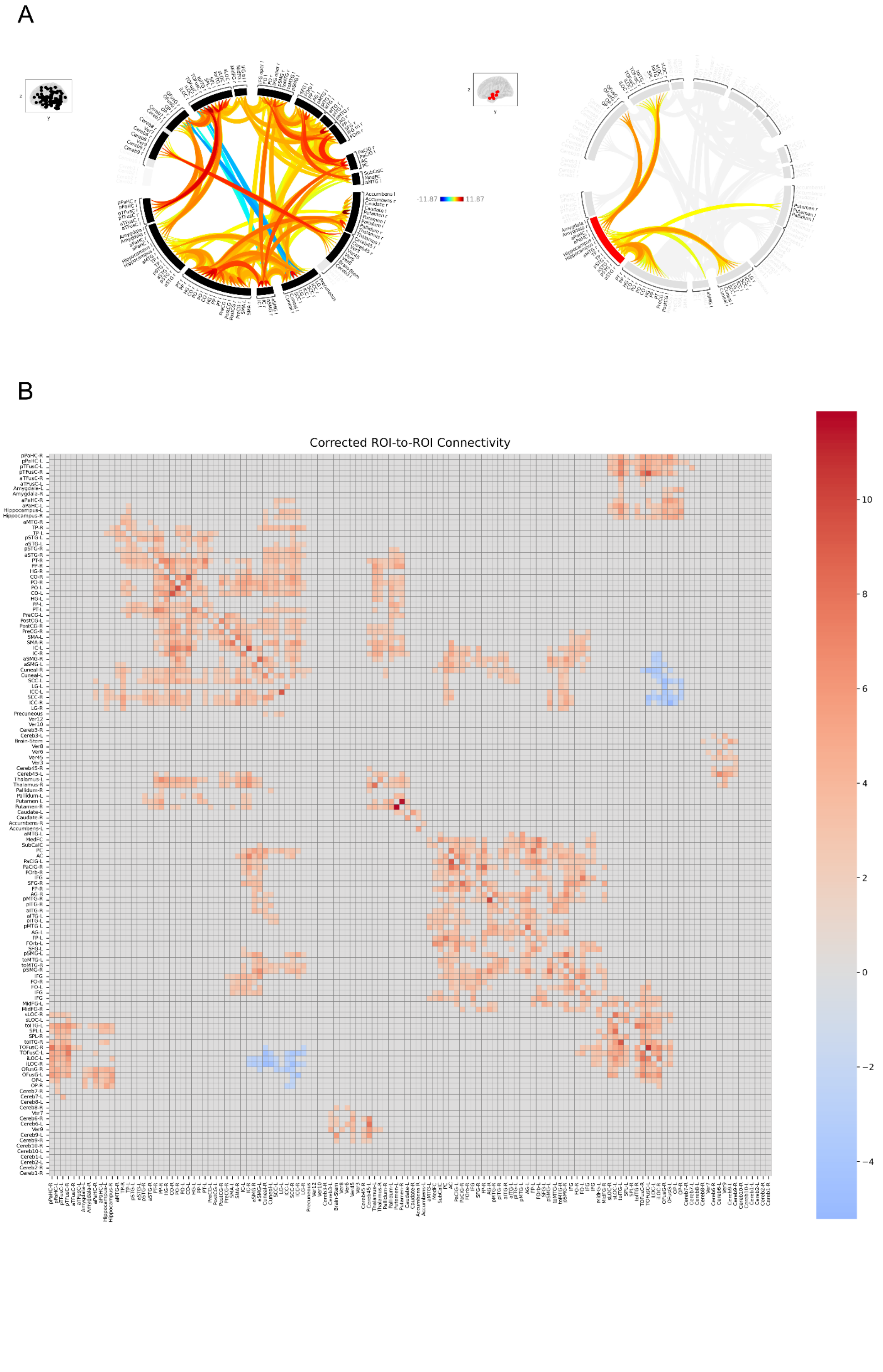

**Figure S4. Task-based ROI-to-ROI functional connectivity during object-location learning (N = 19).** Group-level ROI-to-ROI connectivity matrix (19 participants) derived from generalized psychophysiological interaction (gPPI) analysis for the contrast *learning* > *control*, based on Harvard-Oxford-defined regions of interest (ROIs). Statistical maps were thresholded at $p < 0.001$ with cluster-level family-wise error correction (FWEc) at $p < 0.05$. (A) Connectome rings illustrating ROI-to-ROI functional connectivity, left panel (whole-brain) and right panel (selected OLM-relevant ROI to ROI) (B) Heatmap view for ROI-to-ROI connectivity matrix. The colour scale indicates *t*-values, with warm colours denoting stronger connectivity during learning relative to control, and cool colours denoting the reverse pattern.

**Figure S5:**

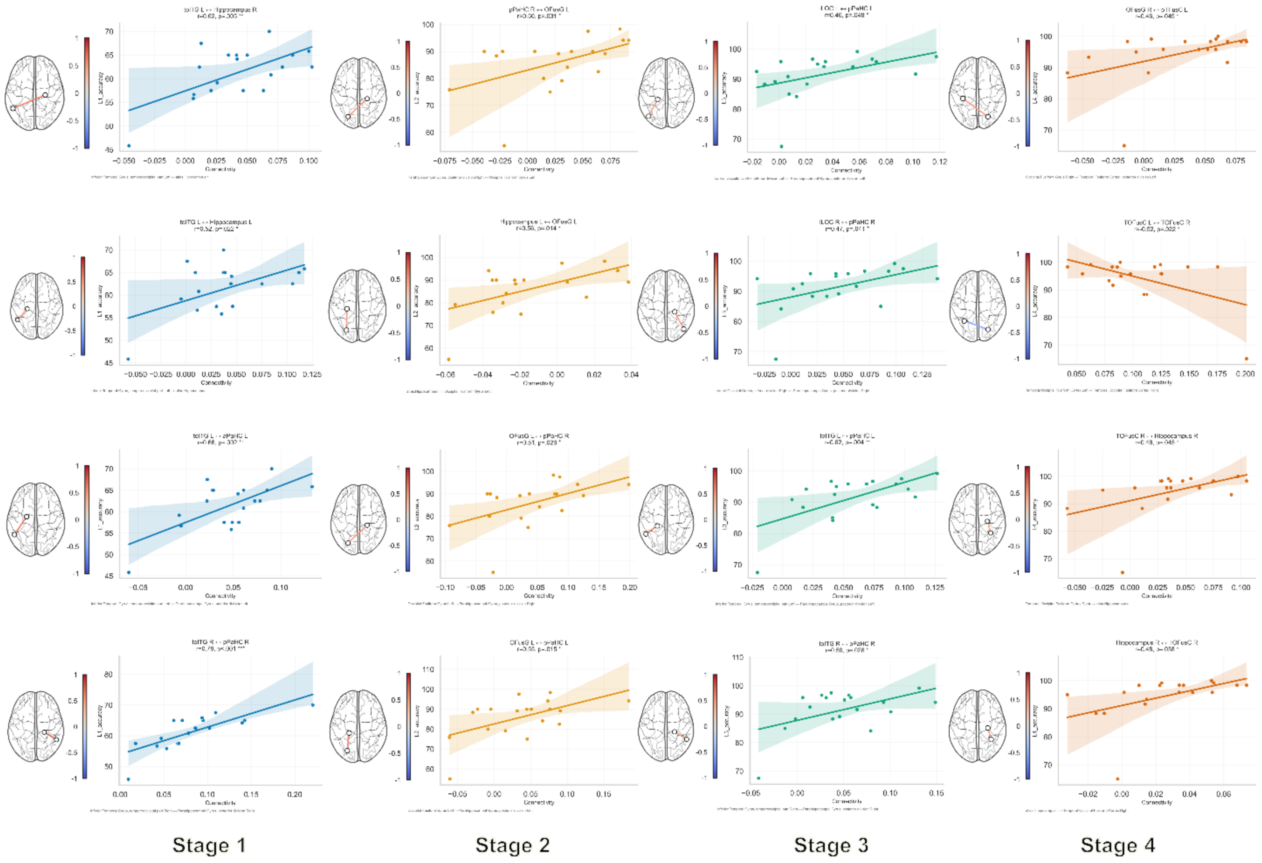

**Figure S5. Stage-specific across-subjects connectivity–behavioral correlations during Object-location learning**

Columns represent learning Stages 1-4, and subplots within each column depict ROI-to-ROI gPPI connectivity-accuracy correlations at the corresponding stage within the a priori OLM-relevant network. For each stage, connectivity estimates were derived from stage-specific gPPI regressors and correlated with behavioral accuracy across participants (Pearson’s r). Positive associations indicate that stronger functional connectivity at a given learning stage was associated with higher performance.

**Figure S6:**

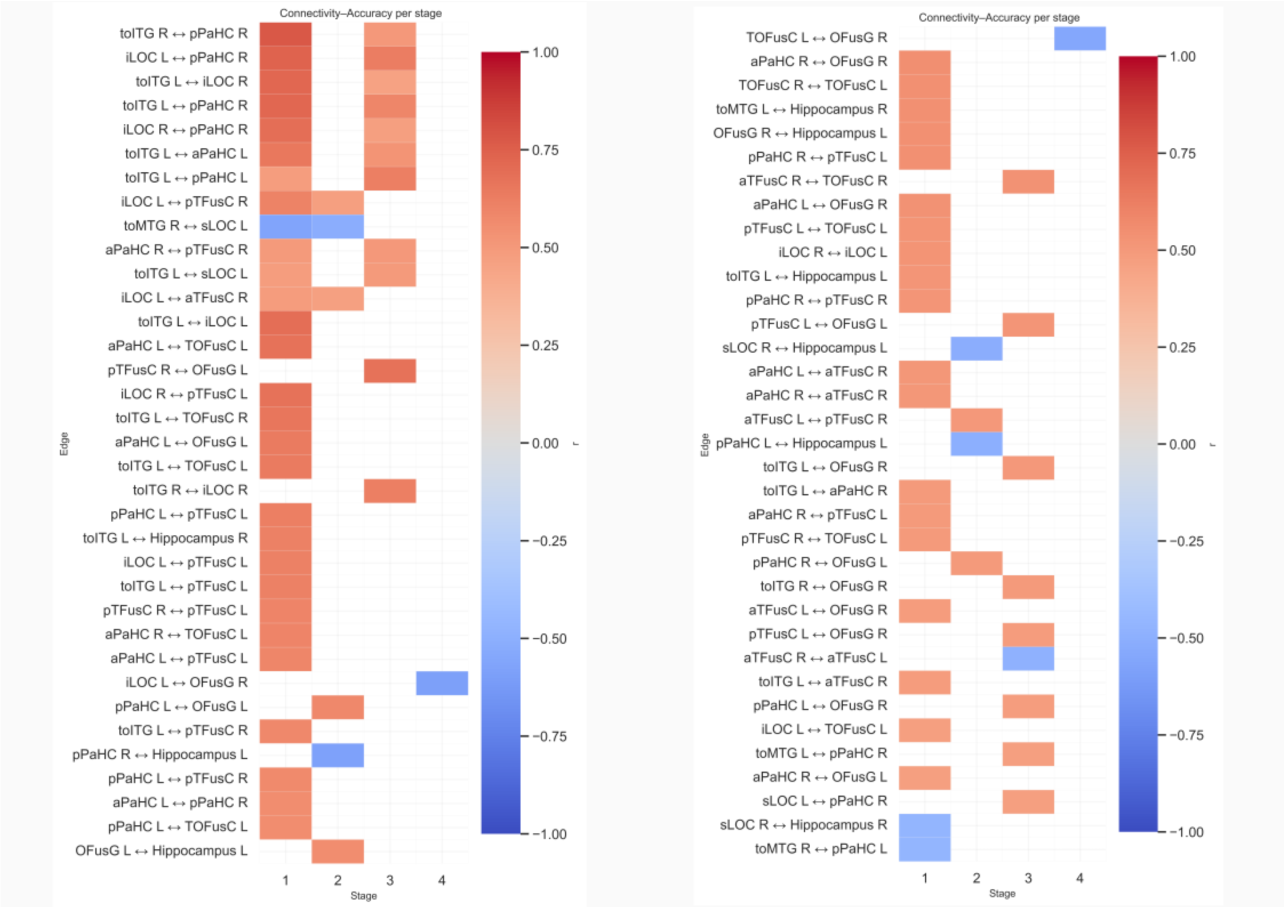

**Figure S6. Stage-specific connectivity–behavior correlations in object-location learning.** Each row is an ROI-to-ROI functional connectivity edge (Harvard-Oxford atlas ROIs from the object-location memory network). Each column is a learning stage (1-4). Color reflects the across-subject Pearson correlation (r) between connectivity strength for that edge and behavioral accuracy at that same stage. Warmer/red cells indicate edges where stronger connectivity is associated with higher accuracy (positive r). Cooler/blue cells indicate edges where stronger connectivity is associated with lower accuracy (negative r). Only edges that showed an uncorrected correlation with behavior ($p < 0.05$) in at least one stage are displayed. Blank cells indicate that the edge was not significant at that stage. The left and right panels show two non-overlapping sets of edges for readability (ranked by effect size magnitude).

**Figure S7:**

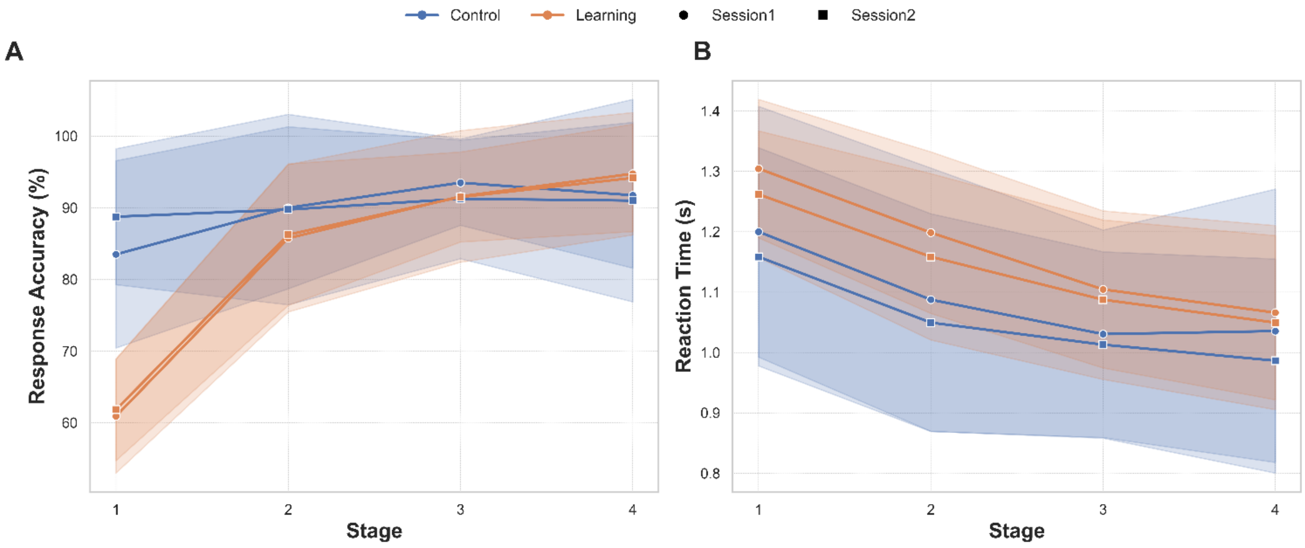

**Figure S7. Behavioral Performance in the Object-Location Memory Task**

(A) Accuracy (%) and (B) reaction time (seconds) for the learning and control tasks across four learning stages (1-4). Results are shown separately for the first fMRI session (circle markers) and the second fMRI session (square markers) in 20 participants. Orange lines represent the learning task, and blue lines represent the control task. Shaded areas indicate standard deviation (SD).

**Table S1: Functionally-connected brain regions to seed: Right Hippocampus in learning>control**

| ROI | Voxels | Percentage | Peak statistics | MNI coordinates (mm) |
|---|---|---|---|---|
| IFG tri l (Inferior Frontal Gyrus, pars triangularis Left) | 200 | 31% | 8.6 | (-52,+30,+4) |
| Hippocampus r | 400 | 57% | 8.2 | (+26,-18,-16) |
| Putamen l | 327 | 38% | 8.1 | (-26,-2,+0) |
| IFG oper l (Inferior Frontal Gyrus, pars opercularis Left) | 43 | 6% | 7.8 | (-44,+8,+26) |
| Putamen r | 146 | 18% | 7.6 | (+30,-4,-2) |
| PreCG l (Precentral Gyrus Left) | 130 | 3% | 7.5 | (-42,+4,+30) |
| MidFG l (Middle Frontal Gyrus Left) | 111 | 4% | 7.5 | (-44,+8,+36) |
| FP l (Frontal Pole Left) | 33 | 0% | 7.4 | (-50,+38,+6) |
| toITG l (Inferior Temporal Gyrus, temporooccipital part Left) | 224 | 32% | 7.0 | (-48,-56,-12) |
| OP r (Occipital Pole Right) | 277 | 11% | 6.5 | (+18,-92,+26) |
| OFusG l (Occipital Fusiform Gyrus Left) | 127 | 14% | 6.4 | (-22,-78,-12) |
| SPL l (Superior Parietal Lobule Left) | 84 | 6% | 6.4 | (-30,-44,+50) |
| sLOC r (Lateral Occipital Cortex, superior division Right) | 105 | 2% | 6.3 | (+22,-86,+26) |
| Amygdala r | 150 | 44% | 6.3 | (+24,-6,-16) |
| OP l (Occipital Pole Left) | 436 | 17% | 5.9 | (-14,-96,+12) |
| IC r (Insular Cortex Right) | 61 | 5% | 5.6 | (+40,+10,-12) |
| toMTG l (Middle Temporal Gyrus, temporooccipital part Left) | 103 | 12% | 5.5 | (-58,-52,-4) |
| OFusG r (Occipital Fusiform Gyrus Right) | 152 | 17% | 5.5 | (+26,-76,-6) |
| aPaHC l (Parahippocampal Gyrus, anterior division Left) | 31 | 5% | 5.4 | (-26,-10,-34) |
| TP r (Temporal Pole Right) | 131 | 5% | 5.4 | (+46,+10,-20) |
| iLOC r (Lateral Occipital Cortex, inferior division Right) | 55 | 3% | 5.3 | (+38,-80,-4) |
| Pallidum l | 36 | 12% | 5.2 | (-22,-2,-2) |
| aTFusC l (Temporal Fusiform Cortex, anterior division Left) | 91 | 29% | 5.2 | (-32,-4,-40) |
| TP l (Temporal Pole Left) | 83 | 4% | 5.1 | (-44,+12,-22) |
| toITG r (Inferior Temporal Gyrus, temporooccipital part Right) | 84 | 11% | 5.0 | (+50,-54,-10) |
| toMTG r (Middle Temporal Gyrus, temporooccipital part Right) | 33 | 3% | 4.9 | (+58,-52,-6) |
| ICC l (Intracalcarine Cortex Left) | 53 | 8% | 4.6 | (-8,-86,+2) |
| iLOC l (Lateral Occipital Cortex, inferior division Left) | 30 | 1% | 4.6 | (-30,-88,+4) |

**Table S2: Functionally-connected brain regions to seed: Left Hippocampus in learning>control**

| ROI | Voxels | Percentage | Peak statistics | MNI coordinates (mm) |
|---|---|---|---|---|
| Hippocampus l | 441 | 58% | 8.9 | (-26,-20,-16) |
| toMTG l (Middle Temporal Gyrus, temporooccipital part Left) | 287 | 33% | 7.6 | (-60,-52,-4) |
| toITG l (Inferior Temporal Gyrus, temporooccipital part Left) | 325 | 47% | 7.1 | (-50,-56,-12) |
| IFG tri l (Inferior Frontal Gyrus, pars triangularis Left) | 345 | 53% | 6.9 | (-50,+30,+8) |
| Amygdala l | 121 | 37% | 6.7 | (-22,-8,-16) |
| FOrb l (Frontal Orbital Cortex Left) | 107 | 6% | 6.6 | (-42,+30,-8) |
| FP l (Frontal Pole Left) | 88 | 1% | 6.4 | (-48,+38,+2) |
| OP r (Occipital Pole Right) | 656 | 26% | 6.4 | (+20,-94,+16) |
| TP r (Temporal Pole Right) | 112 | 5% | 6.3 | (+44,+10,-22) |
| OFusG r (Occipital Fusiform Gyrus Right) | 409 | 46% | 6.2 | (+26,-76,-10) |
| sLOC r (Lateral Occipital Cortex, superior division Right) | 103 | 2% | 6.1 | (+22,-86,+26) |
| toMTG r (Middle Temporal Gyrus, temporooccipital part Right) | 243 | 21% | 6.1 | (+60,-50,-4) |
| OP l (Occipital Pole Left) | 271 | 10% | 5.9 | (-10,-96,+10) |
| LG l (Lingual Gyrus Left) | 232 | 15% | 5.8 | (-10,-78,-8) |
| FP l (Frontal Pole Left) | 172 | 2% | 5.8 | (-10,+52,+40) |
| PreCG l (Precentral Gyrus Left) | 51 | 1% | 5.7 | (-40,+4,+30) |
| MidFG l (Middle Frontal Gyrus Left) | 58 | 2% | 5.7 | (-42,+8,+36) |
| toITG r (Inferior Temporal Gyrus, temporooccipital part Right) | 88 | 11% | 5.6 | (+56,-52,-12) |
| TOFusC l (Temporal Occipital Fusiform Cortex Left) | 46 | 7% | 5.4 | (-40,-54,-14) |
| ICC l (Intracalcarine Cortex Left) | 83 | 13% | 5.4 | (-6,-84,+2) |
| OFusG l (Occipital Fusiform Gyrus Left) | 63 | 7% | 5.3 | (-20,-76,-12) |
| MidFG l (Middle Frontal Gyrus Left) | 77 | 3% | 5.2 | (-50,+32,+24) |
| iLOC r (Lateral Occipital Cortex, inferior division Right) | 36 | 2% | 5.1 | (+38,-80,-8) |
| ICC r (Intracalcarine Cortex Right) | 50 | 7% | 4.6 | (+10,-84,+6) |

**Table S3: Functionally-connected brain regions to seed: Right Parahippocampus in learning>control contrast**

| ROI | Voxels | Percentage | Peak statistics | MNI coordinates (mm) |
|---|---|---|---|---|
| pPaHC r (Parahippocampal Gyrus, posterior division Right) | 201 | 63% | 13.6 | (+22,-32,-16) |
| LG r (Lingual Gyrus Right) | 91 | 5% | 10.3 | (+24,-42,-10) |
| pTFusC r (Temporal Fusiform Cortex, posterior division Right) | 91 | 13% | 8.6 | (+30,-36,-20) |
| Cereb45 r (Cerebelum 4 5 Right) | 37 | 6% | 7.9 | (+18,-36,-20) |
| Cereb3 r (Cerebelum 3 Right) | 31 | 17% | 7.2 | (+14,-32,-18) |
| TOFusC r (Temporal Occipital Fusiform Cortex Right) | 620 | 76% | 7.0 | (+34,-50,-16) |
| OFusG l (Occipital Fusiform Gyrus Left) | 502 | 54% | 6.9 | (-28,-76,-14) |
| toITG r (Inferior Temporal Gyrus, temporooccipital part Right) | 155 | 20% | 6.7 | (+48,-54,-12) |
| sLOC l (Lateral Occipital Cortex, superior division Left) | 822 | 17% | 6.6 | (-30,-76,+26) |
| TOFusC l (Temporal Occipital Fusiform Cortex Left) | 374 | 57% | 6.5 | (-32,-54,-14) |
| OFusG r (Occipital Fusiform Gyrus Right) | 242 | 27% | 6.4 | (+34,-68,-14) |
| pTFusC l (Temporal Fusiform Cortex, posterior division Left) | 76 | 9% | 6.3 | (-32,-42,-16) |
| iLOC r (Lateral Occipital Cortex, inferior division Right) | 322 | 16% | 6.3 | (+40,-76,-6) |
| SFG l (Superior Frontal Gyrus Left) | 178 | 6% | 6.3 | (-6,+12,+64) |
| SMA L(Juxtapositional Lobule Cortex -formerly Supplementary Motor Cortex- Left) | 78 | 12% | 6.2 | (-4,+6,+62) |
| LG l (Lingual Gyrus Left) | 46 | 3% | 6.2 | (-26,-48,-8) |
| toITG l (Inferior Temporal Gyrus, temporooccipital part Left) | 267 | 38% | 6.1 | (-48,-56,-14) |
| PreCG l (Precentral Gyrus Left) | 156 | 4% | 6.0 | (-44,+2,+40) |
| Cereb6 r (Cerebelum 6 Right) | 113 | 7% | 5.6 | (+32,-60,-22) |
| iLOC l (Lateral Occipital Cortex, inferior division Left) | 514 | 25% | 5.5 | (-40,-80,-6) |
| sLOC r (Lateral Occipital Cortex, superior division Right) | 288 | 6% | 5.4 | (+30,-68,+34) |
| PaCiG l (Paracingulate Gyrus Left) | 47 | 4% | 4.9 | (-4,+16,+48) |
| SPL l (Superior Parietal Lobule Left) | 89 | 6% | 4.9 | (-28,-48,+48) |
| MidFG l (Middle Frontal Gyrus Left) | 36 | 1% | 4.8 | (-44,+6,+38) |
| OP l (Occipital Pole Left) | 123 | 5% | 4.8 | (-28,-92,+0) |

Functionally-connected brain regions to seed: Right Parahippocampus in control>learning contrast

| ROI | Voxels | Percentage | Peak statistics | MNI coordinates (mm) |
|---|---|---|---|---|
| PC (Cingulate Gyrus, posterior division) | 254 | 11% | 7.0 | (+4,-48,+30) |

| | | | | |
|---|---|---|---|---|
| Precuneous (Precuneous Cortex) | 101 | 2% | 5.0 | (+0,-50,+38) |
| FP l (Frontal Pole Left) | 97 | 1% | 5.2 | (-8,+60,+2) |
| FP r (Frontal Pole Right) | 66 | 1% | 4.8 | (+6,+58,+4) |
| PaCiG l (Paracingulate Gyrus Left) | 77 | 6% | 4.7 | (-6,+40,-6) |
| SubCalC (Subcallosal Cortex) | 38 | 3% | 4.9 | (+2,+30,-8) |

**Table S4: Functionally-connected brain regions to seed: Left Parahippocampus in learning>control contrast**

| ROI | Voxels | Percentage | Peak statistics | MNI coordinates (mm) |
|---|---|---|---|---|
| IFG oper l (Inferior Frontal Gyrus, pars opercularis Left) | 397 | 52% | 9.2 | (-48,+16,+20) |
| PreCG l (Precentral Gyrus Left) | 337 | 8% | 8.8 | (-46,+2,+38) |
| pTFusC l (Temporal Fusiform Cortex, posterior division Left) | 217 | 25% | 8.5 | (-32,-36,-20) |
| SPL l (Superior Parietal Lobule Left) | 586 | 40% | 8.4 | (-30,-50,+50) |
| MidFG l (Middle Frontal Gyrus Left) | 706 | 24% | 8.4 | (-44,+18,+34) |
| TOFusC l (Temporal Occipital Fusiform Cortex Left) | 402 | 62% | 8.0 | (-32,-54,-14) |
| sLOC l (Lateral Occipital Cortex, superior division Left) | 1017 | 20% | 7.8 | (-28,-74,+34) |
| pPaHC l (Parahippocampal Gyrus, posterior division Left) | 220 | 56% | 7.8 | (-24,-34,-16) |
| toITG l (Inferior Temporal Gyrus, temporooccipital part Left) | 428 | 61% | 7.5 | (-50,-56,-14) |
| OFusG l (Occipital Fusiform Gyrus Left) | 457 | 49% | 7.4 | (-30,-72,-14) |
| IFG tri l (Inferior Frontal Gyrus, pars triangularis Left) | 414 | 64% | 7.4 | (-50,+30,+12) |
| SFG l (Superior Frontal Gyrus Left) | 155 | 5% | 7.2 | (-6,+14,+60) |
| iLOC l (Lateral Occipital Cortex, inferior division Left) | 464 | 23% | 6.2 | (-40,-78,-10) |
| SFG r (Superior Frontal Gyrus Right) | 46 | 2% | 6.2 | (+2,+16,+56) |
| LG r (Lingual Gyrus Right) | 117 | 7% | 6.1 | (+28,-44,-8) |
| iLOC r (Lateral Occipital Cortex, inferior division Right) | 295 | 14% | 6.1 | (+40,-78,-10) |
| LG l (Lingual Gyrus Left) | 173 | 11% | 6.1 | (-24,-54,-8) |
| OFusG r (Occipital Fusiform Gyrus Right) | 264 | 30% | 6.1 | (+34,-68,-12) |
| toITG r (Inferior Temporal Gyrus, temporooccipital part Right) | 106 | 14% | 6.1 | (+48,-56,-12) |
| IFG tri r (Inferior Frontal Gyrus, pars triangularis Right) | 123 | 22% | 6.0 | (+50,+30,+16) |
| PostCG l (Postcentral Gyrus Left) | 79 | 2% | 6.0 | (-40,-36,+44) |
| SPL r (Superior Parietal Lobule Right) | 183 | 12% | 6.0 | (+30,-50,+48) |
| SMA L(Juxtapositional Lobule Cortex -formerly Supplementary Motor Cortex- Left) | 74 | 12% | 6.0 | (-4,+6,+60) |
| Cereb6 l (Cerebelum 6 Left) | 46 | 4% | 5.9 | (-18,-74,-18) |

| | | | | |
|---|---|---|---|---|
| TOFusC r (Temporal Occipital Fusiform Cortex Right) | 406 | 50% | 5.8 | (+34,-52,-14) |
| SFG l (Superior Frontal Gyrus Left) | 124 | 4% | 5.7 | (-24,+6,+60) |
| MidFG r (Middle Frontal Gyrus Right) | 133 | 5% | 5.6 | (+50,+26,+28) |
| TP l (Temporal Pole Left) | 94 | 4% | 5.6 | (-34,+6,-40) |
| aSMG l (Supramarginal Gyrus, anterior division Left) | 56 | 6% | 5.5 | (-42,-38,+42) |
| sLOC r (Lateral Occipital Cortex, superior division Right) | 382 | 8% | 5.4 | (+26,-66,+40) |
| pTFusC r (Temporal Fusiform Cortex, posterior division Right) | 51 | 7% | 5.4 | (+30,-36,-18) |
| aTFusC l (Temporal Fusiform Cortex, anterior division Left) | 72 | 23% | 5.3 | (-32,-4,-40) |
| pSMG l (Supramarginal Gyrus, posterior division Left) | 30 | 3% | 5.0 | (-38,-46,+42) |
| Cereb6 r (Cerebelum 6 Right) | 65 | 4% | 4.8 | (+30,-62,-22) |
| pPaHC r (Parahippocampal Gyrus, posterior division Right) | 49 | 15% | 4.8 | (+26,-36,-14) |
| OP l (Occipital Pole Left) | 68 | 3% | 4.8 | (-32,-92,+4) |

Functionally-connected brain regions to seed: Left Parahippocampus in control>learning contrast

| ROI | Voxels | Percentage | Peak statistics | MNI coordinates (mm) |
|---|---|---|---|---|
| PaCiG l (Paracingulate Gyrus Left) | 91 | 7% | 7.0 | (-6,+16,+48) |
| PaCiG r (Paracingulate Gyrus Right) | 34 | 2% | 6.2 | (+4,+16,+50) |
| PC (Cingulate Gyrus, posterior division) | 68 | 3% | 5.6 | (+6,-46,+30) |
| FP l (Frontal Pole Left) | 54 | 1% | 5.5 | (-6,+56,+2) |
| PaCiG r (Paracingulate Gyrus Right) | 151 | 11% | 5.4 | (+6,+52,+10) |
| FP r (Frontal Pole Right) | 68 | 1% | 5.4 | (+6,+58,+6) |
| PaCiG l (Paracingulate Gyrus Left) | 138 | 11% | 5.2 | (-6,+50,+0) |
| AC (Cingulate Gyrus, anterior division) | 98 | 4% | 4.8 | (-2,+38,+0) |

**Table S5: Functionally-connected brain regions to seed: Right Lateral Occipital Cortex (LOC) in learning>control contrast**

| ROI | Voxels | Percentage | Peak statistics | MNI coordinates (mm) |
|---|---|---|---|---|
| iLOC r (Lateral Occipital Cortex, inferior division Right) | 1304 | 64% | 11.0 | (+44,-78,-4) |
| OFusG r (Occipital Fusiform Gyrus Right) | 213 | 24% | 10.1 | (+36,-74,-14) |
| TOFusC r (Temporal Occipital Fusiform Cortex Right) | 519 | 64% | 8.6 | (+36,-50,-16) |
| iLOC l (Lateral Occipital Cortex, inferior division Left) | 627 | 31% | 8.6 | (-38,-82,-8) |
| toITG r (Inferior Temporal Gyrus, temporooccipital part Right) | 137 | 18% | 8.0 | (+48,-54,-12) |

| | | | | |
|---|---|---|---|---|
| pTFusC r (Temporal Fusiform Cortex, posterior division Right) | 163 | 23% | 7.6 | (+34,-32,-20) |
| OFusG l (Occipital Fusiform Gyrus Left) | 221 | 24% | 7.5 | (-32,-78,-14) |
| LG r (Lingual Gyrus Right) | 96 | 6% | 7.3 | (+28,-42,-8) |
| sLOC l (Lateral Occipital Cortex, superior division Left) | 1252 | 25% | 7.1 | (-28,-76,+36) |
| pPaHC r (Parahippocampal Gyrus, posterior division Right) | 132 | 41% | 6.9 | (+28,-32,-18) |
| pTFusC l (Temporal Fusiform Cortex, posterior division Left) | 152 | 18% | 6.4 | (-32,-40,-18) |
| TOFusC l (Temporal Occipital Fusiform Cortex Left) | 190 | 29% | 6.2 | (-32,-52,-16) |
| sLOC r (Lateral Occipital Cortex, superior division Right) | 407 | 8% | 6.2 | (+42,-76,+26) |
| Cereb1 l (Cerebelum Crus1 Left) | 37 | 2% | 6.2 | (-34,-84,-22) |
| SFG l (Superior Frontal Gyrus Left) | 33 | 1% | 6.0 | (-6,+24,+50) |
| pPaHC l (Parahippocampal Gyrus, posterior division Left) | 83 | 21% | 5.8 | (-28,-36,-16) |
| Precuneous (Precuneous Cortex) | 58 | 1% | 5.5 | (+18,-54,+22) |
| Cereb6 r (Cerebelum 6 Right) | 77 | 5% | 5.4 | (+36,-60,-24) |
| Cereb1 r (Cerebelum Crus1 Right) | 54 | 2% | 5.2 | (+40,-62,-26) |

Functionally-connected brain regions to seed: Right Lateral Occipital Cortex (LOC) in control>learning contrast

| ROI | Voxels | Percentage | Peak statistics | MNI coordinates (mm) |
|---|---|---|---|---|
| OP r (Occipital Pole Right) | 349 | 14% | 7.8 | (+30,-94,-6) |
| CO r (Central Opercular Cortex Right) | 110 | 13% | 7.1 | (+44,+6,+6) |
| IC r (Insular Cortex Right) | 148 | 11% | 6.8 | (+38,+6,+4) |
| OP l (Occipital Pole Left) | 327 | 12% | 6.7 | (-28,-94,-6) |
| PC (Cingulate Gyrus, posterior division) | 83 | 3% | 6.3 | (+8,-18,+42) |
| FO r (Frontal Operculum Cortex Right) | 40 | 13% | 6.1 | (+42,+12,+8) |
| AC (Cingulate Gyrus, anterior division) | 182 | 7% | 6.0 | (+0,-2,+42) |
| IFG oper r (Inferior Frontal Gyrus, pars opercularis Right) | 91 | 13% | 5.7 | (+54,+14,+6) |
| Cuneal l (Cuneal Cortex Left) | 176 | 34% | 5.6 | (-6,-84,+24) |
| PaCiG l (Paracingulate Gyrus Left) | 70 | 5% | 5.6 | (-6,+24,+42) |
| Cuneal r (Cuneal Cortex Right) | 249 | 39% | 5.5 | (+6,-80,+26) |
| SCC r (Supracalcarine Cortex Right) | 44 | 31% | 5.3 | (+2,-80,+14) |
| PO l (Parietal Operculum Cortex Left) | 71 | 13% | 5.2 | (-52,-30,+22) |
| OP r (Occipital Pole Right) | 109 | 4% | 5.1 | (+6,-90,+22) |
| pSMG r (Supramarginal Gyrus, posterior division Right) | 143 | 12% | 5.1 | (+60,-38,+44) |
| aSMG r (Supramarginal Gyrus, anterior division Right) | 80 | 10% | 5.1 | (+60,-32,+40) |
| PostCG l (Postcentral Gyrus Left) | 69 | 2% | 4.8 | (-60,-18,+20) |
| CO l (Central Opercular Cortex Left) | 32 | 3% | 4.7 | (-54,-18,+20) |

**Table S6: Functionally-connected brain regions to seed: Left Lateral Occipital Cortex (LOC) in learning>control contrast**

| ROI | Voxels | Percentage | Peak statistics | MNI coordinates (mm) |
|---|---|---|---|---|
| iLOC l (Lateral Occipital Cortex, inferior division Left) | 1299 | 64% | 11.3 | (-42,-78,-4) |
| iLOC r (Lateral Occipital Cortex, inferior division Right) | 1037 | 51% | 9.3 | (+44,-76,-4) |
| TOFusC r (Temporal Occipital Fusiform Cortex Right) | 722 | 89% | 9.1 | (+34,-50,-16) |
| OFusG r (Occipital Fusiform Gyrus Right) | 131 | 15% | 8.7 | (+38,-66,-14) |
| TOFusC l (Temporal Occipital Fusiform Cortex Left) | 517 | 79% | 8.6 | (-34,-54,-16) |
| sLOC l (Lateral Occipital Cortex, superior division Left) | 2123 | 43% | 8.3 | (-28,-74,+40) |
| pTFusC r (Temporal Fusiform Cortex, posterior division Right) | 213 | 30% | 8.3 | (+34,-32,-22) |
| toITG r (Inferior Temporal Gyrus, temporooccipital part Right) | 195 | 25% | 8.2 | (+50,-54,-14) |
| LG r (Lingual Gyrus Right) | 140 | 8% | 8.0 | (+26,-44,-8) |
| pPaHC r (Parahippocampal Gyrus, posterior division Right) | 174 | 55% | 8.0 | (+26,-32,-18) |
| OFusG l (Occipital Fusiform Gyrus Left) | 82 | 9% | 6.8 | (-36,-70,-12) |
| pTFusC l (Temporal Fusiform Cortex, posterior division Left) | 192 | 22% | 6.8 | (-32,-40,-18) |
| toITG l (Inferior Temporal Gyrus, temporooccipital part Left) | 121 | 17% | 6.7 | (-48,-60,-14) |
| Hippocampus l | 42 | 6% | 6.2 | (-28,-36,-8) |
| MidFG l (Middle Frontal Gyrus Left) | 136 | 5% | 6.2 | (-44,+22,+32) |
| sLOC r (Lateral Occipital Cortex, superior division Right) | 976 | 20% | 6.1 | (+36,-74,+32) |
| Cereb6 r (Cerebelum 6 Right) | 105 | 7% | 6.0 | (+34,-56,-24) |
| pPaHC l (Parahippocampal Gyrus, posterior division Left) | 116 | 30% | 5.9 | (-26,-36,-16) |
| LG l (Lingual Gyrus Left) | 82 | 5% | 5.8 | (-24,-46,-10) |
| Cereb1 r (Cerebelum Crus1 Right) | 81 | 3% | 5.7 | (+42,-58,-30) |
| SPL l (Superior Parietal Lobule Left) | 285 | 20% | 5.3 | (-24,-54,+58) |
| MidFG l (Middle Frontal Gyrus Left) | 84 | 3% | 5.2 | (-30,+8,+56) |
| Precuneous (Precuneous Cortex) | 99 | 2% | 5.2 | (+18,-56,+22) |
| SFG l (Superior Frontal Gyrus Left) | 52 | 2% | 5.1 | (-24,+6,+56) |
| Precuneous (Precuneous Cortex) | 138 | 2% | 4.9 | (-10,-60,+46) |

Functionally-connected brain regions to seed: Left Lateral Occipital Cortex (LOC) in control>learning contrast

| ROI | Voxels | Percentage | Peak statistics | MNI coordinates (mm) |
|---|---|---|---|---|
| IC r (Insular Cortex Right) | 88 | 7% | 5.8 | (+36,+4,+6) |
| OP r (Occipital Pole Right) | 32 | 1% | 5.5 | (+38,-92,+0) |
| CO l (Central Opercular Cortex Left) | 44 | 4% | 5.3 | (-54,-20,+18) |
| OP l (Occipital Pole Left) | 86 | 3% | 5.2 | (-34,-90,+6) |
| PO l (Parietal Operculum Cortex Left) | 63 | 11% | 5.0 | (-54,-28,+20) |
| Cuneal r (Cuneal Cortex Right) | 100 | 16% | 4.9 | (+4,-84,+24) |
| Cuneal l (Cuneal Cortex Left) | 99 | 19% | 4.5 | (-6,-84,+24) |
| OP r (Occipital Pole Right) | 113 | 5% | 4.8 | (+4,-92,+20) |

**Table S7: Functionally-connected brain regions to seed: Right Middle Temporal Gyrus (Temporo-occipital part) in learning>control contrast**

| ROI | Voxels | Percentage | Peak statistics | MNI coordinates (mm) |
|---|---|---|---|---|
| toMTG r (Middle Temporal Gyrus, temporooccipital part Right) | 731 | 63% | 11.8 | (+60,-48,+2) |
| pSMG r (Supramarginal Gyrus, posterior division Right) | 531 | 43% | 6.7 | (+56,-42,+36) |
| AG r (Angular Gyrus Right) | 435 | 30% | 6.3 | (+56,-50,+24) |
| pMTG r (Middle Temporal Gyrus, posterior division Right) | 195 | 14% | 5.7 | (+60,-32,-6) |
| FP r (Frontal Pole Right) | 193 | 2% | 5.7 | (+14,+48,+40) |
| pSMG l (Supramarginal Gyrus, posterior division Left) | 231 | 22% | 5.7 | (-52,-46,+48) |
| AC (Cingulate Gyrus, anterior division) | 209 | 8% | 5.7 | (+0,+34,+12) |
| PaCiG l (Paracingulate Gyrus Left) | 90 | 7% | 5.4 | (-10,+44,+16) |
| AG l (Angular Gyrus Left) | 70 | 7% | 5.2 | (-46,-54,+52) |
| aSMG l (Supramarginal Gyrus, anterior division Left) | 65 | 7% | 5.1 | (-54,-38,+48) |
| pITG r (Inferior Temporal Gyrus, posterior division Right) | 93 | 10% | 5.0 | (+58,-22,-28) |
| aSMG r (Supramarginal Gyrus, anterior division Right) | 85 | 11% | 4.9 | (+56,-32,+46) |
| toITG r (Inferior Temporal Gyrus, temporooccipital part Right) | 33 | 4% | 4.5 | (+60,-44,-18) |

**Table S8: Functionally-connected brain regions to seed: Left Middle Temporal Gyrus (Temporo-occipital part) in learning>control contrast**

| ROI | Voxels | Percentage | Peak statistics | MNI coordinates (mm) |
|---|---|---|---|---|
| toMTG l (Middle Temporal Gyrus, temporooccipital part Left) | 621 | 72% | 12.7 | (-58,-54,+2) |

| | | | | |
|---|---|---|---|---|
| pSMG l (Supramarginal Gyrus, posterior division Left) | 74 | 7% | 7.7 | (-56,-46,+10) |
| iLOC l (Lateral Occipital Cortex, inferior division Left) | 40 | 2% | 7.1 | (-58,-64,+6) |
| pITG r (Inferior Temporal Gyrus, posterior division Right) | 193 | 20% | 6.0 | (+54,-18,-30) |
| AG l (Angular Gyrus Left) | 169 | 18% | 5.9 | (-56,-56,+18) |
| PC (Cingulate Gyrus, posterior division) | 193 | 8% | 5.6 | (-4,-30,+42) |
| pMTG r (Middle Temporal Gyrus, posterior division Right) | 93 | 7% | 5.4 | (+60,-16,-24) |
| pSMG r (Supramarginal Gyrus, posterior division Right) | 87 | 7% | 5.3 | (+64,-40,+28) |
| SubCalC (Subcallosal Cortex) | 177 | 16% | 5.0 | (-4,+22,-10) |
| AG r (Angular Gyrus Right) | 67 | 5% | 4.7 | (+56,-56,+18) |
| toMTG r (Middle Temporal Gyrus, temporooccipital part Right) | 41 | 4% | 4.5 | (+54,-54,+8) |
| sLOC l (Lateral Occipital Cortex, superior division Left) | 40 | 1% | 4.5 | (-54,-62,+20) |
| sLOC r (Lateral Occipital Cortex, superior division Right) | 37 | 1% | 4.5 | (+56,-62,+22) |

**Table S9: Functionally-connected brain regions to seed: right inferior temporal gyrus, temporo-occipital in learning>control contrast**

| ROI | Voxels | Percentage | Peak statistics | MNI coordinates (mm) |
|---|---|---|---|---|
| toITG r (Inferior Temporal Gyrus, temporooccipital part Right) | 552 | 71% | 11.1 | (+52,-50,-16) |
| TOFusC r (Temporal Occipital Fusiform Cortex Right) | 501 | 61% | 10.5 | (+38,-48,-16) |
| toITG l (Inferior Temporal Gyrus, temporooccipital part Left) | 554 | 79% | 9.8 | (-50,-54,-16) |
| SubCalC (Subcallosal Cortex) | 119 | 11% | 8.4 | (-6,+18,-18) |
| TOFusC l (Temporal Occipital Fusiform Cortex Left) | 180 | 28% | 8.3 | (-36,-48,-18) |
| iLOC r (Lateral Occipital Cortex, inferior division Right) | 93 | 5% | 8.2 | (+46,-64,-12) |
| iLOC l (Lateral Occipital Cortex, inferior division Left) | 334 | 16% | 7.6 | (-48,-68,-10) |
| OFusG r (Occipital Fusiform Gyrus Right) | 35 | 4% | 7.4 | (+40,-64,-14) |
| pTFusC r (Temporal Fusiform Cortex, posterior division Right) | 178 | 25% | 7.3 | (+34,-34,-20) |
| pTFusC l (Temporal Fusiform Cortex, posterior division Left) | 259 | 30% | 6.5 | (-34,-38,-20) |
| sLOC r (Lateral Occipital Cortex, superior division Right) | 482 | 10% | 6.2 | (+36,-66,+38) |
| pPaHC l (Parahippocampal Gyrus, posterior division Left) | 42 | 11% | 6.1 | (-30,-34,-14) |

| | | | | |
|---|---|---|---|---|
| Hippocampus l | 76 | 10% | 5.7 | (-28,-36,-6) |
| pPaHC r (Parahippocampal Gyrus, posterior division Right) | 86 | 27% | 5.5 | (+26,-32,-18) |
| sLOC l (Lateral Occipital Cortex, superior division Left) | 264 | 5% | 5.5 | (-34,-76,+32) |
| Cereb1 r (Cerebelum Crus1 Right) | 51 | 2% | 5.5 | (+46,-48,-28) |
| toMTG l (Middle Temporal Gyrus, temporooccipital part Left) | 90 | 10% | 5.4 | (-56,-58,-6) |
| toMTG r (Middle Temporal Gyrus, temporooccipital part Right) | 30 | 3% | 5.3 | (+56,-50,-8) |
| Precuneous (Precuneous Cortex) | 48 | 1% | 5.1 | (+16,-54,+18) |
| LG r (Lingual Gyrus Right) | 85 | 5% | 5.0 | (+28,-42,-8) |

**Table S10: Functionally-connected brain regions to seed: left inferior temporal gyrus, temporo-occipital part in learning>control contrast**

| ROI | Voxels | Percentage | Peak statistics | MNI coordinates (mm) |
|---|---|---|---|---|
| toITG l (Inferior Temporal Gyrus, temporooccipital part Left) | 552 | 79% | 14.8 | (-50,-54,-16) |
| iLOC l (Lateral Occipital Cortex, inferior division Left) | 369 | 18% | 12.4 | (-50,-66,-6) |
| toMTG l (Middle Temporal Gyrus, temporooccipital part Left) | 89 | 10% | 10.1 | (-52,-60,-2) |
| toITG r (Inferior Temporal Gyrus, temporooccipital part Right) | 524 | 67% | 9.1 | (+52,-52,-16) |
| sLOC r (Lateral Occipital Cortex, superior division Right) | 1116 | 23% | 8.7 | (+38,-70,+28) |
| sLOC l (Lateral Occipital Cortex, superior division Left) | 1029 | 21% | 8.2 | (-32,-76,+30) |
| TOFusC r (Temporal Occipital Fusiform Cortex Right) | 651 | 80% | 8.1 | (+36,-50,-16) |
| TOFusC l (Temporal Occipital Fusiform Cortex Left) | 311 | 48% | 7.5 | (-34,-50,-14) |
| iLOC r (Lateral Occipital Cortex, inferior division Right) | 438 | 21% | 7.1 | (+46,-68,-4) |
| pTFusC l (Temporal Fusiform Cortex, posterior division Left) | 287 | 33% | 6.9 | (-34,-38,-20) |
| LG r (Lingual Gyrus Right) | 145 | 8% | 6.7 | (+26,-44,-10) |
| pTFusC r (Temporal Fusiform Cortex, posterior division Right) | 199 | 28% | 6.7 | (+34,-32,-20) |
| pPaHC r (Parahippocampal Gyrus, posterior division Right) | 174 | 55% | 6.7 | (+26,-32,-18) |
| Hippocampus l | 190 | 25% | 6.6 | (-28,-28,-14) |
| LG l (Lingual Gyrus Left) | 68 | 4% | 6.6 | (-26,-44,-8) |
| pPaHC l (Parahippocampal Gyrus, posterior division Left) | 190 | 49% | 6.5 | (-26,-34,-16) |
| OFusG r (Occipital Fusiform Gyrus Right) | 39 | 4% | 6.3 | (+40,-64,-14) |

| | | | | |
|---|---|---|---|---|
| Cereb1 r (Cerebelum Crus1 Right) | 73 | 3% | 5.9 | (+48,-50,-28) |
| toMTG r (Middle Temporal Gyrus, temporooccipital part Right) | 55 | 5% | 5.7 | (+54,-54,-6) |
| Hippocampus r | 31 | 4% | 5.7 | (+30,-32,-10) |

**Table S11: Functionally-connected brain regions to activation-based seed: right temporal-occipital fusiform cortex in learning>control contrast**

| ROI | Voxels | Percentage | Peak statistics | MNI coordinates (mm) |
|---|---|---|---|---|
| TOFusC r (Temporal Occipital Fusiform Cortex Right) | 638 | 78% | 21.7 | (+34,-50,-16) |
| LG r (Lingual Gyrus Right) | 178 | 10% | 14.8 | (+26,-44,-8) |
| pTFusC r (Temporal Fusiform Cortex, posterior division Right) | 142 | 20% | 12.1 | (+32,-34,-20) |
| TOFusC l (Temporal Occipital Fusiform Cortex Left) | 556 | 85% | 10.7 | (-32,-54,-16) |
| pPaHC r (Parahippocampal Gyrus, posterior division Right) | 208 | 65% | 10.1 | (+24,-32,-16) |
| pTFusC l (Temporal Fusiform Cortex, posterior division Left) | 201 | 23% | 9.8 | (-32,-40,-20) |
| OFusG l (Occipital Fusiform Gyrus Left) | 528 | 57% | 9.3 | (-30,-76,-14) |
| LG l (Lingual Gyrus Left) | 97 | 6% | 8.8 | (-26,-48,-8) |
| pPaHC l (Parahippocampal Gyrus, posterior division Left) | 90 | 23% | 7.8 | (-26,-36,-16) |
| OFusG r (Occipital Fusiform Gyrus Right) | 246 | 28% | 7.7 | (+34,-70,-12) |
| iLOC l (Lateral Occipital Cortex, inferior division Left) | 704 | 34% | 7.7 | (-40,-82,-8) |
| Cereb6 r (Cerebelum 6 Right) | 133 | 9% | 7.0 | (+32,-56,-24) |
| iLOC r (Lateral Occipital Cortex, inferior division Right) | 721 | 35% | 6.9 | (+40,-80,-4) |
| toITG r (Inferior Temporal Gyrus, temporooccipital part Right) | 171 | 22% | 6.9 | (+50,-54,-12) |
| sLOC r (Lateral Occipital Cortex, superior division Right) | 458 | 10% | 6.8 | (+36,-74,+26) |
| OP r (Occipital Pole Right) | 156 | 6% | 6.3 | (+34,-92,-4) |
| PaCiG l (Paracingulate Gyrus Left) | 104 | 8% | 6.3 | (-6,+20,+44) |
| sLOC l (Lateral Occipital Cortex, superior division Left) | 699 | 14% | 6.1 | (-34,-80,+24) |
| SFG l (Superior Frontal Gyrus Left) | 66 | 2% | 5.9 | (-4,+18,+52) |
| Cereb45 r (Cerebelum 4 5 Right) | 36 | 6% | 5.7 | (+22,-38,-22) |
| Cereb45 l (Cerebelum 4 5 Left) | 38 | 4% | 5.6 | (-24,-44,-20) |
| OP l (Occipital Pole Left) | 300 | 11% | 5.6 | (-28,-94,-4) |
| Cereb6 l (Cerebelum 6 Left) | 77 | 6% | 5.3 | (-30,-56,-28) |
| Precuneous (Precuneous Cortex) | 54 | 1% | 5.3 | (+18,-54,+20) |
| toITG l (Inferior Temporal Gyrus, temporooccipital part Left) | 145 | 21% | 5.1 | (-48,-56,-14) |
| PaCiG r (Paracingulate Gyrus Right) | 38 | 3% | 4.9 | (+4,+20,+44) |
| Thalamus r | 36 | 3% | 4.7 | (+18,-30,+0) |

| | | | | |
|---|---|---|---|---|
| Cereb1 l (Cerebelum Crus1 Left) | 44 | 2% | 4.7 | (-32,-76,-22) |

Functionally-connected brain regions to activation-based seed: right temporal-occipital fusiform cortex in control>learning contrast

| ROI | Voxels | Percentage | Peak statistics | MNI coordinates (mm) |
|---|---|---|---|---|
| PreCG r (Precentral Gyrus Right) | 29 | 1% | 11.01 | (+56,-6,+24) |
| PostCG r (Postcentral Gyrus Right) | 29 | 1% | 7.58 | (+56,-8,+24) |
| CO r (Central Opercular Cortex Right) | 19 | 2% | 5.41 | (+52,-6,+14) |

**Table S12: Functionally-connected brain regions to activation-based seed: left temporal-occipital fusiform cortex in learning>control contrast**

| ROI | Voxels | Percentage | Peak statistics | MNI coordinates (mm) |
|---|---|---|---|---|
| TOFusC l (Temporal Occipital Fusiform Cortex Left) | 595 | 91% | 26.0 | (-34,-54,-16) |
| Cereb6 l (Cerebelum 6 Left) | 226 | 17% | 11.9 | (-32,-54,-24) |
| OFusG l (Occipital Fusiform Gyrus Left) | 231 | 25% | 11.1 | (-32,-74,-14) |
| sLOC r (Lateral Occipital Cortex, superior division Right) | 888 | 18% | 9.7 | (+30,-70,+44) |
| iLOC l (Lateral Occipital Cortex, inferior division Left) | 395 | 19% | 7.3 | (-38,-82,-8) |
| PaCiG l (Paracingulate Gyrus Left) | 163 | 12% | 7.3 | (-6,+20,+44) |
| TOFusC r (Temporal Occipital Fusiform Cortex Right) | 314 | 39% | 7.2 | (+34,-46,-16) |
| SFG l (Superior Frontal Gyrus Left) | 81 | 3% | 7.0 | (-4,+18,+52) |
| pTFusC r (Temporal Fusiform Cortex, posterior division Right) | 168 | 23% | 7.0 | (+32,-34,-22) |
| pPaHC r (Parahippocampal Gyrus, posterior division Right) | 154 | 48% | 6.7 | (+26,-32,-18) |
| pTFusC l (Temporal Fusiform Cortex, posterior division Left) | 206 | 24% | 6.6 | (-32,-40,-20) |
| toITG r (Inferior Temporal Gyrus, temporooccipital part Right) | 132 | 17% | 6.5 | (+48,-54,-12) |
| sLOC l (Lateral Occipital Cortex, superior division Left) | 252 | 5% | 6.5 | (-18,-66,+50) |
| toITG l (Inferior Temporal Gyrus, temporooccipital part Left) | 205 | 29% | 6.3 | (-48,-56,-14) |
| iLOC r (Lateral Occipital Cortex, inferior division Right) | 141 | 7% | 6.2 | (+48,-64,-10) |
| iLOC r (Lateral Occipital Cortex, inferior division Right) | 196 | 10% | 6.1 | (+36,-86,-4) |
| OP l (Occipital Pole Left) | 217 | 8% | 6.0 | (-32,-94,-4) |
| pPaHC l (Parahippocampal Gyrus, posterior division Left) | 38 | 10% | 5.9 | (-26,-38,-14) |
| sLOC l (Lateral Occipital Cortex, superior division Left) | 473 | 10% | 5.6 | (-34,-82,+26) |

| | | | | |
|---|---|---|---|---|
| PaCiG r (Paracingulate Gyrus Right) | 66 | 5% | 5.4 | (+4,+20,+46) |
| OP r (Occipital Pole Right) | 35 | 1% | 5.4 | (+34,-92,-2) |
| MidFG l (Middle Frontal Gyrus Left) | 137 | 5% | 5.3 | (-42,+22,+30) |
| Cereb45 l (Cerebelum 4 5 Left) | 31 | 3% | 5.2 | (-24,-44,-20) |
| Cereb6 r (Cerebelum 6 Right) | 88 | 6% | 5.2 | (+34,-60,-22) |
| OFusG r (Occipital Fusiform Gyrus Right) | 62 | 7% | 5.1 | (+38,-64,-16) |
| IFG oper l (Inferior Frontal Gyrus, pars opercularis Left) | 32 | 4% | 5.1 | (-40,+14,+26) |
| SMA L(Juxtapositional Lobule Cortex - formerly Supplementary Motor Cortex- Left) | 30 | 5% | 4.6 | (-6,+6,+58) |

Functionally-connected brain regions to activation-based seed: left temporal-occipital fusiform cortex in control>learning contrast

| ROI | Voxels | Percentage | Peak statistics | MNI coordinates (mm) |
|---|---|---|---|---|
| LG r (Lingual Gyrus Right) | 94 | 5% | 6.8 | (+26,-42,-10) |
| Cuneal r (Cuneal Cortex Right) | 147 | 23% | 5.9 | (+6,-80,+24) |
| LG l (Lingual Gyrus Left) | 44 | 3% | 6.0 | (-24,-46,-10) |
| ICC r (Intracalcarine Cortex Right) | 266 | 35% | 5.8 | (+10,-78,+6) |
| LG r (Lingual Gyrus Right) | 331 | 19% | 5.6 | (+10,-76,-6) |
| SCC r (Supracalcarine Cortex Right) | 52 | 36% | 5.6 | (+4,-80,+12) |

**Table S13: Functionally-connected brain regions to activation-based seed: right temporal fusiform cortex, posterior division in learning>control contrast**

| ROI | Voxels | Percentage | Peak statistics | MNI coordinates (mm) |
|---|---|---|---|---|
| pTFusC r (Temporal Fusiform Cortex, posterior division Right) | 173 | 24% | 16.8 | (+32,-34,-20) |
| TOFusC r (Temporal Occipital Fusiform Cortex Right) | 494 | 61% | 11.7 | (+34,-52,-16) |
| Cereb45 r (Cerebelum 4 5 Right) | 49 | 8% | 10.0 | (+24,-38,-24) |
| pPaHC r (Parahippocampal Gyrus, posterior division Right) | 117 | 37% | 7.9 | (+24,-34,-16) |
| TOFusC l (Temporal Occipital Fusiform Cortex Left) | 450 | 69% | 7.7 | (-34,-54,-16) |
| sLOC l (Lateral Occipital Cortex, superior division Left) | 1001 | 20% | 7.6 | (-28,-74,+36) |
| sLOC r (Lateral Occipital Cortex, superior division Right) | 837 | 17% | 7.5 | (+32,-68,+36) |
| LG r (Lingual Gyrus Right) | 97 | 6% | 7.3 | (+28,-42,-8) |
| pTFusC l (Temporal Fusiform Cortex, posterior division Left) | 153 | 18% | 6.8 | (-34,-40,-18) |
| iLOC r (Lateral Occipital Cortex, inferior division Right) | 124 | 6% | 6.8 | (+48,-64,-12) |
| Precuneous (Precuneous Cortex) | 59 | 1% | 6.8 | (+20,-56,+22) |
| SPL l (Superior Parietal Lobule Left) | 45 | 3% | 6.8 | (-22,-56,+54) |
| PreCG l (Precentral Gyrus Left) | 39 | 1% | 6.6 | (-4,-22,+50) |

| | | | | |
|---|---|---|---|---|
| PaCiG l (Paracingulate Gyrus Left) | 78 | 6% | 6.5 | (-6,+22,+44) |
| toITG l (Inferior Temporal Gyrus, temporooccipital part Left) | 350 | 50% | 6.5 | (-50,-56,-16) |
| LG l (Lingual Gyrus Left) | 61 | 4% | 6.4 | (-26,-46,-8) |
| toITG r (Inferior Temporal Gyrus, temporooccipital part Right) | 192 | 25% | 6.4 | (+50,-54,-12) |
| iLOC l (Lateral Occipital Cortex, inferior division Left) | 371 | 18% | 6.4 | (-42,-76,-10) |
| OFusG r (Occipital Fusiform Gyrus Right) | 94 | 11% | 6.2 | (+38,-64,-14) |
| MidFG l (Middle Frontal Gyrus Left) | 131 | 4% | 6.2 | (-42,+20,+28) |
| Cereb6 r (Cerebelum 6 Right) | 54 | 3% | 6.0 | (+36,-56,-24) |
| iLOC r (Lateral Occipital Cortex, inferior division Right) | 253 | 12% | 5.8 | (+38,-82,+0) |
| IFG tri l (Inferior Frontal Gyrus, pars triangularis Left) | 58 | 9% | 5.8 | (-46,+28,+18) |
| MidFG l (Middle Frontal Gyrus Left) | 54 | 2% | 5.6 | (-28,+8,+52) |
| SFG l (Superior Frontal Gyrus Left) | 69 | 2% | 5.6 | (-22,+8,+54) |
| OFusG l (Occipital Fusiform Gyrus Left) | 122 | 13% | 5.4 | (-38,-72,-14) |
| OP l (Occipital Pole Left) | 59 | 2% | 4.9 | (-34,-94,-8) |
| Cereb1 r (Cerebelum Crus1 Right) | 34 | 1% | 4.8 | (+42,-58,-26) |
| SPL r (Superior Parietal Lobule Right) | 43 | 3% | 4.6 | (+20,-54,+58) |
| IFG oper l (Inferior Frontal Gyrus, pars opercularis Left) | 49 | 6% | 4.4 | (-42,+12,+26) |

Functionally-connected brain regions to activation-based seed: right temporal fusiform cortex, posterior division in control>learning contrast

| ROI | Voxels | Percentage | Peak statistics | MNI coordinates (mm) |
|---|---|---|---|---|
| PC (Cingulate Gyrus, posterior division) | 112 | 5% | 6.7 | (+2,-22,+42) |
| PreCG l (Precentral Gyrus Left) | 234 | 5% | 6.7 | (-48,+0,+44) |
| PC (Cingulate Gyrus, posterior division) | 186 | 8% | 6.0 | (+4,-46,+28) |
| AG l (Angular Gyrus Left) | 71 | 7% | 5.3 | (-48,-56,+34) |
| sLOC l (Lateral Occipital Cortex, superior division Left) | 112 | 2% | 4.8 | (-48,-62,+44) |

**Table S14: Functionally-connected brain regions to activation-based seed: Left temporal fusiform cortex, posterior division in learning>control contrast**

| ROI | Voxels | Percentage | Peak statistics | MNI coordinates (mm) |
|---|---|---|---|---|
| pTFusC l (Temporal Fusiform Cortex, posterior division Left) | 211 | 25% | 16.1 | (-32,-38,-20) |
| pPaHC l (Parahippocampal Gyrus, posterior division Left) | 105 | 27% | 15.1 | (-26,-36,-16) |
| PreCG l (Precentral Gyrus Left) | 308 | 7% | 8.9 | (-44,+2,+38) |
| MidFG l (Middle Frontal Gyrus Left) | 799 | 27% | 8.8 | (-44,+14,+36) |
| OFusG r (Occipital Fusiform Gyrus Right) | 336 | 38% | 8.3 | (+34,-70,-12) |
| iLOC r (Lateral Occipital Cortex, inferior division Right) | 724 | 35% | 8.3 | (+40,-78,-6) |
| TOFusC r (Temporal Occipital Fusiform Cortex Right) | 555 | 68% | 8.0 | (+34,-52,-16) |
| toITG l (Inferior Temporal Gyrus, temporooccipital part Left) | 422 | 60% | 8.0 | (-50,-56,-14) |
| sLOC l (Lateral Occipital Cortex, superior division Left) | 1205 | 24% | 8.0 | (-26,-74,+36) |
| TOFusC l (Temporal Occipital Fusiform Cortex Left) | 401 | 62% | 7.9 | (-32,-54,-14) |
| iLOC l (Lateral Occipital Cortex, inferior division Left) | 676 | 33% | 7.9 | (-42,-78,-8) |
| SPL l (Superior Parietal Lobule Left) | 478 | 33% | 7.8 | (-30,-50,+50) |
| pTFusC r (Temporal Fusiform Cortex, posterior division Right) | 74 | 10% | 7.8 | (+30,-36,-18) |
| toITG r (Inferior Temporal Gyrus, temporooccipital part Right) | 151 | 19% | 7.5 | (+48,-56,-12) |
| OFusG l (Occipital Fusiform Gyrus Left) | 417 | 45% | 7.4 | (-30,-74,-14) |
| IFG oper l (Inferior Frontal Gyrus, pars opercularis Left) | 148 | 19% | 7.4 | (-44,+14,+24) |
| Cereb6 r (Cerebelum 6 Right) | 234 | 15% | 7.2 | (+30,-62,-26) |
| pPaHC r (Parahippocampal Gyrus, posterior division Right) | 67 | 21% | 7.1 | (+26,-34,-14) |
| sLOC r (Lateral Occipital Cortex, superior division Right) | 971 | 20% | 7.1 | (+32,-68,+40) |
| LG l (Lingual Gyrus Left) | 136 | 9% | 7.0 | (-26,-48,-6) |
| LG r (Lingual Gyrus Right) | 153 | 9% | 6.9 | (+26,-44,-8) |
| Cereb1 r (Cerebelum Crus1 Right) | 143 | 6% | 6.7 | (+36,-60,-30) |
| SPL r (Superior Parietal Lobule Right) | 158 | 11% | 6.1 | (+30,-48,+46) |
| IFG tri l (Inferior Frontal Gyrus, pars triangularis Left) | 41 | 6% | 6.1 | (-44,+26,+20) |
| PaCiG l (Paracingulate Gyrus Left) | 92 | 7% | 6.0 | (-6,+16,+48) |
| SFG l (Superior Frontal Gyrus Left) | 121 | 4% | 5.9 | (-24,+6,+60) |
| SFG l (Superior Frontal Gyrus Left) | 154 | 5% | 5.9 | (-6,+14,+58) |
| OP l (Occipital Pole Left) | 190 | 7% | 5.8 | (-30,-94,+0) |
| Cereb8 r (Cerebelum 8 Right) | 88 | 4% | 5.5 | (+18,-68,-40) |
| PaCiG r (Paracingulate Gyrus Right) | 31 | 2% | 5.5 | (+4,+16,+50) |

| | | | | |
|---|---|---|---|---|
| SFG r (Superior Frontal Gyrus Right) | 38 | 1% | 5.3 | (+2,+16,+56) |
| OP r (Occipital Pole Right) | 111 | 4% | 5.3 | (+36,-92,+0) |
| PostCG l (Postcentral Gyrus Left) | 34 | 1% | 4.9 | (-38,-36,+44) |
| SMA L(Juxtapositional Lobule Cortex - formerly Supplementary Motor Cortex- Left) | 38 | 6% | 4.9 | (-4,+6,+62) |
| Precuneous (Precuneous Cortex) | 40 | 1% | 4.7 | (+12,-58,+52) |

Functionally-connected brain regions to activation-based seed: Left temporal fusiform cortex, posterior division in control>learning contrast

| ROI | Voxels | Percentage | Peak statistics | MNI coordinates (mm) |
|---|---|---|---|---|
| FP l (Frontal Pole Left) | 108 | 2% | 8.0 | (-6,+58,+2) |
| AC (Cingulate Gyrus, anterior division) | 312 | 12% | 7.4 | (+0,+38,+0) |
| FP r (Frontal Pole Right) | 268 | 3% | 7.1 | (+8,+56,+14) |
| PaCiG r (Paracingulate Gyrus Right) | 471 | 35% | 6.7 | (+8,+48,+4) |
| PaCiG l (Paracingulate Gyrus Left) | 345 | 26% | 6.6 | (-6,+46,-2) |
| PC (Cingulate Gyrus, posterior division) | 220 | 9% | 6.2 | (+2,-48,+26) |
| SubCalC (Subcallosal Cortex) | 54 | 5% | 5.7 | (+2,+30,-6) |
| MedFC (Frontal Medial Cortex) | 261 | 27% | 5.6 | (+0,+48,-14) |
| Precuneous (Precuneous Cortex) | 60 | 1% | 5.5 | (-4,-60,+22) |
| SFG r (Superior Frontal Gyrus Right) | 47 | 2% | 5.4 | (+6,+54,+26) |

**Table S15: Functionally-connected brain regions to activation-based seed: right lateral occipital cortex in learning>control contrast**

| ROI | Voxels | Percentage | Peak statistics | MNI coordinates (mm) |
|---|---|---|---|---|
| iLOC r (Lateral Occipital Cortex, inferior division Right) | 1015 | 50% | 14.2 | (+42,-78,-4) |
| LG r (Lingual Gyrus Right) | 113 | 7% | 9.4 | (+28,-42,-8) |
| toITG r (Inferior Temporal Gyrus, temporooccipital part Right) | 144 | 18% | 9.3 | (+48,-56,-12) |
| sLOC l (Lateral Occipital Cortex, superior division Left) | 839 | 17% | 9.0 | (-34,-80,+26) |
| TOFusC r (Temporal Occipital Fusiform Cortex Right) | 421 | 52% | 9.0 | (+34,-50,-16) |
| Precuneous (Precuneous Cortex) | 142 | 3% | 8.6 | (+16,-56,+22) |
| sLOC r (Lateral Occipital Cortex, superior division Right) | 242 | 5% | 8.1 | (+38,-80,+22) |
| Precuneous (Precuneous Cortex) | 99 | 2% | 7.8 | (-16,-60,+22) |
| pPaHC r (Parahippocampal Gyrus, posterior division Right) | 109 | 34% | 6.9 | (+28,-32,-18) |
| pTFusC r (Temporal Fusiform Cortex, posterior division Right) | 130 | 18% | 6.7 | (+32,-34,-20) |
| OFusG r (Occipital Fusiform Gyrus Right) | 86 | 10% | 6.1 | (+40,-68,-14) |

| | | | | |
|---|---|---|---|---|
| iLOC l (Lateral Occipital Cortex, inferior division Left) | 581 | 28% | 6.1 | (-40,-78,-2) |
| CO r (Central Opercular Cortex Right) | 113 | 13% | 5.9 | (+44,+6,+6) |
| pTFusC l (Temporal Fusiform Cortex, posterior division Left) | 74 | 9% | 5.9 | (-30,-40,-16) |
| TOFusC l (Temporal Occipital Fusiform Cortex Left) | 115 | 18% | 5.8 | (-28,-48,-14) |
| pPaHC l (Parahippocampal Gyrus, posterior division Left) | 54 | 14% | 5.7 | (-28,-36,-14) |
| LG l (Lingual Gyrus Left) | 31 | 2% | 5.7 | (-28,-44,-8) |
| Cereb6 r (Cerebelum 6 Right) | 56 | 4% | 5.3 | (+36,-60,-24) |
| Cereb1 r (Cerebelum Crus1 Right) | 37 | 1% | 5.1 | (+40,-60,-26) |
| Hippocampus l | 48 | 6% | 5.0 | (-30,-34,-8) |
| OP r (Occipital Pole Right) | 104 | 4% | 4.9 | (+32,-92,-10) |

Functionally-connected brain regions to activation-based seed: right lateral occipital cortex in control>learning contrast

| ROI | Voxels | Percentage | Peak statistics | MNI coordinates (mm) |
|---|---|---|---|---|
| PostCG l (Postcentral Gyrus Left) | 269 | 7% | 8.0 | (-58,-18,+26) |
| aSMG l (Supramarginal Gyrus, anterior division Left) | 200 | 21% | 7.7 | (-60,-28,+30) |
| PO l (Parietal Operculum Cortex Left) | 78 | 14% | 6.7 | (-54,-30,+22) |
| CO l (Central Opercular Cortex Left) | 61 | 6% | 6.6 | (-56,-18,+18) |
| FO r (Frontal Operculum Cortex Right) | 98 | 31% | 6.6 | (+44,+14,+6) |
| OP r (Occipital Pole Right) | 291 | 12% | 6.1 | (+10,-92,+18) |
| FP r (Frontal Pole Right) | 289 | 4% | 5.9 | (+22,+50,+32) |
| IC r (Insular Cortex Right) | 97 | 7% | 5.7 | (+36,+6,+6) |
| IFG oper r (Inferior Frontal Gyrus, pars opercularis Right) | 102 | 15% | 5.6 | (+54,+12,+6) |
| PostCG r (Postcentral Gyrus Right) | 173 | 5% | 5.6 | (+54,-18,+34) |
| aSMG r (Supramarginal Gyrus, anterior division Right) | 241 | 30% | 5.4 | (+60,-26,+34) |
| pSMG r (Supramarginal Gyrus, posterior division Right) | 164 | 13% | 5.3 | (+58,-40,+42) |
| toMTG l (Middle Temporal Gyrus, temporooccipital part Left) | 66 | 8% | 5.3 | (-62,-52,+8) |
| SCC r (Supracalcarine Cortex Right) | 42 | 29% | 5.2 | (+2,-80,+12) |
| Cuneal r (Cuneal Cortex Right) | 30 | 5% | 5.0 | (+8,-84,+20) |
| ICC r (Intracalcarine Cortex Right) | 39 | 5% | 4.9 | (+8,-84,+12) |
| AG r (Angular Gyrus Right) | 32 | 2% | 4.5 | (+56,-48,+42) |
| OP l (Occipital Pole Left) | 46 | 2% | 4.5 | (-4,-98,+4) |

**Table S16: Functionally-connected brain regions to activation-based seed: left lateral occipital cortex in learning>control contrast**

| ROI | Voxels | Percentage | Peak statistics | MNI coordinates (mm) |
|---|---|---|---|---|
| sLOC l (Lateral Occipital Cortex, superior division Left) | 1955 | 39% | 14.8 | (-24,-76,+38) |
| OP l (Occipital Pole Left) | 192 | 7% | 10.0 | (-30,-92,+4) |
| Precuneous (Precuneous Cortex) | 311 | 6% | 7.6 | (-16,-62,+30) |
| Precuneous (Precuneous Cortex) | 106 | 2% | 6.4 | (+18,-56,+20) |
| TOFusC r (Temporal Occipital Fusiform Cortex Right) | 247 | 30% | 6.4 | (+34,-48,-14) |
| iLOC r (Lateral Occipital Cortex, inferior division Right) | 655 | 32% | 6.2 | (+40,-78,-4) |
| sLOC r (Lateral Occipital Cortex, superior division Right) | 379 | 8% | 6.2 | (+34,-72,+34) |
| toITG r (Inferior Temporal Gyrus, temporooccipital part Right) | 69 | 9% | 6.1 | (+46,-56,-10) |
| pTFusC r (Temporal Fusiform Cortex, posterior division Right) | 130 | 18% | 6.0 | (+32,-34,-20) |
| TOFusC l (Temporal Occipital Fusiform Cortex Left) | 175 | 27% | 5.7 | (-30,-54,-14) |
| iLOC l (Lateral Occipital Cortex, inferior division Left) | 177 | 9% | 5.7 | (-34,-86,+0) |
| OP r (Occipital Pole Right) | 81 | 3% | 5.7 | (+34,-92,+2) |
| pPaHC r (Parahippocampal Gyrus, posterior division Right) | 79 | 25% | 5.6 | (+28,-32,-18) |
| LG r (Lingual Gyrus Right) | 48 | 3% | 5.5 | (+30,-42,-8) |
| PreCG l (Precentral Gyrus Left) | 46 | 1% | 5.4 | (-40,+2,+34) |
| PaCiG r (Paracingulate Gyrus Right) | 44 | 3% | 5.2 | (+6,+38,+28) |
| SPL l (Superior Parietal Lobule Left) | 83 | 6% | 4.9 | (-18,-56,+58) |
| pPaHC l (Parahippocampal Gyrus, posterior division Left) | 30 | 8% | 4.8 | (-28,-38,-12) |
| pTFusC l (Temporal Fusiform Cortex, posterior division Left) | 55 | 6% | 4.8 | (-30,-40,-14) |
| iLOC l (Lateral Occipital Cortex, inferior division Left) | 55 | 3% | 4.6 | (-44,-68,-8) |
| OFusG l (Occipital Fusiform Gyrus Left) | 58 | 6% | 4.4 | (-36,-70,-14) |

Functionally-connected brain regions to activation-based seed: left lateral occipital cortex in control>learning contrast

| ROI | Voxels | Percentage | Peak statistics | MNI coordinates (mm) |
|---|---|---|---|---|
| FP r (Frontal Pole Right) | 299 | 4% | 6.4 | (+24,+48,+28) |
| pSMG r (Supramarginal Gyrus, posterior division Right) | 264 | 21% | 5.8 | (+58,-40,+42) |
| aSMG r (Supramarginal Gyrus, anterior division Right) | 98 | 12% | 5.0 | (+60,-32,+42) |
| IC r (Insular Cortex Right) | 146 | 11% | 5.0 | (+38,+0,-6) |

**Table S17: Significant Relevant ROI-to-ROI Connections**

| ROI A | ROI B | t-stat | p-FWE |
|---|---|---|---|
| **atlas.TOFusC-R** | atlas.TOFusC-L | 10.590 | 2.1e-05 |
| **atlas.pTFusC-R** | atlas.TOFusC-L | 9.540 | 4.6e-05 |
| **atlas.toITG-R** | atlas.toITG-L | 9.200 | 5.9e-05 |
| **atlas.iLOC-L** | atlas.TOFusC-R | 7.920 | 0.000234 |
| **atlas.pTFusC-R** | atlas.TOFusC-R | 7.710 | 0.000298 |
| **atlas.TOFusC-R** | atlas.pPaHC-R | 7.660 | 0.0003 |
| **atlas.sLOC-R** | atlas.sLOC-L | 7.620 | 0.000304 |
| **atlas.TOFusC-L** | atlas.pTFusC-R | 7.400 | 0.000401 |
| **atlas.toITG-L** | atlas.toITG-R | 7.390 | 0.000401 |
| **atlas.pTFusC-R** | atlas.toITG-L | 7.320 | 0.00041 |
| **atlas.TOFusC-R** | atlas.pTFusC-R | 7.290 | 0.000416 |
| **atlas.TOFusC-L** | atlas.TOFusC-R | 7.200 | 0.00047 |
| **atlas.iLOC-L** | atlas.TOFusC-L | 7.030 | 0.000556 |
| **atlas.pPaHC-L** | atlas.toITG-L | 7.010 | 0.000556 |
| **atlas.iLOC-L** | atlas.iLOC-R | 6.990 | 0.000556 |
| **atlas.OFusG-L** | atlas.TOFusC-R | 6.730 | 0.000826 |
| **atlas.pTFusC-L** | atlas.toITG-L | 6.570 | 0.000964 |
| **atlas.TOFusC-R** | atlas.iLOC-R | 6.500 | 0.001043 |
| **atlas.toITG-L** | atlas.TOFusC-R | 6.470 | 0.001066 |
| **atlas.iLOC-L** | atlas.sLOC-L | 6.150 | 0.001743 |
| **atlas.toITG-L** | atlas.pTFusC-R | 6.010 | 0.002127 |
| **atlas.TOFusC-R** | atlas.toITG-L | 5.950 | 0.002338 |
| **atlas.OFusG-L** | atlas.iLOC-R | 5.940 | 0.002338 |
| **atlas.iLOC-L** | atlas.pTFusC-R | 5.920 | 0.002346 |
| **atlas.pPaHC-R** | atlas.OFusG-L | 5.900 | 0.002458 |
| **atlas.Hippocampus-L** | atlas.toITG-L | 5.860 | 0.002568 |
| **atlas.TOFusC-R** | atlas.pTFusC-L | 5.810 | 0.002629 |
| **atlas.aPaHC-L** | atlas.toITG-L | 5.810 | 0.002629 |
| **atlas.OFusG-L** | atlas.pPaHC-R | 5.780 | 0.002762 |
| **atlas.pTFusC-R** | atlas.iLOC-L | 5.690 | 0.003118 |
| **atlas.toITG-L** | atlas.sLOC-L | 5.650 | 0.003385 |
| **atlas.toITG-R** | atlas.sLOC-L | 5.620 | 0.003488 |
| **atlas.pPaHC-R** | atlas.TOFusC-R | 5.620 | 0.003488 |
| **atlas.aPaHC-R** | atlas.OFusG-L | 5.600 | 0.003495 |
| **atlas.iLOC-R** | atlas.TOFusC-R | 5.580 | 0.003502 |
| **atlas.OFusG-L** | atlas.TOFusC-L | 5.550 | 0.003653 |
| **atlas.OFusG-R** | atlas.TOFusC-R | 5.480 | 0.003957 |
| **atlas.pTFusC-L** | atlas.TOFusC-R | 5.470 | 0.003968 |
| **atlas.toITG-L** | atlas.pTFusC-L | 5.460 | 0.004041 |
| **atlas.toITG-R** | atlas.TOFusC-R | 5.450 | 0.004054 |
| **atlas.toITG-L** | atlas.pPaHC-L | 5.450 | 0.004054 |
| **atlas.iLOC-L** | atlas.pPaHC-R | 5.440 | 0.004098 |
| **atlas.TOFusC-L** | atlas.toITG-L | 5.340 | 0.004571 |
| **atlas.OFusG-L** | atlas.Hippocampus-R | 5.280 | 0.00497 |
| **atlas.OFusG-L** | atlas.aPaHC-R | 5.280 | 0.00497 |
| **atlas.toITG-L** | atlas.TOFusC-L | 5.270 | 0.005014 |
| **atlas.aPaHC-R** | atlas.OFusG-R | 5.190 | 0.005469 |
| **atlas.TOFusC-L** | atlas.iLOC-R | 5.180 | 0.005487 |
| **atlas.TOFusC-L** | atlas.pPaHC-R | 5.130 | 0.005756 |
| **atlas.TOFusC-R** | atlas.iLOC-L | 5.080 | 0.006134 |
| **atlas.sLOC-R** | atlas.TOFusC-L | 5.050 | 0.006441 |
| **atlas.pPaHC-R** | atlas.toITG-L | 5.030 | 0.006528 |
| **atlas.pTFusC-L** | atlas.TOFusC-L | 4.980 | 0.007028 |
| **atlas.pPaHC-L** | atlas.TOFusC-L | 4.950 | 0.007299 |
| **atlas.sLOC-L** | atlas.sLOC-R | 4.910 | 0.007658 |
| **atlas.Hippocampus-L** | atlas.OFusG-R | 4.890 | 0.007812 |
| **atlas.OFusG-L** | atlas.sLOC-R | 4.870 | 0.007962 |
| **atlas.TOFusC-L** | atlas.sLOC-R | 4.860 | 0.007972 |

| | | | |
|---|---|---|---|
| **atlas.toITG-R** | atlas.iLOC-L | 4.850 | 0.008136 |
| **atlas.pTFusC-R** | atlas.iLOC-R | 4.850 | 0.008136 |
| **atlas.OFusG-L** | atlas.pTFusC-R | 4.840 | 0.008228 |
| **atlas.TOFusC-R** | atlas.sLOC-L | 4.820 | 0.008341 |
| **atlas.OFusG-L** | atlas.pPaHC-L | 4.810 | 0.008362 |
| **atlas.OFusG-R** | atlas.aPaHC-R | 4.800 | 0.008513 |
| **atlas.toITG-R** | atlas.TOFusC-L | 4.800 | 0.008536 |
| **atlas.TOFusC-R** | atlas.toITG-R | 4.760 | 0.009 |
| **atlas.iLOC-R** | atlas.TOFusC-L | 4.710 | 0.009646 |
| **atlas.TOFusC-R** | atlas.OFusG-L | 4.700 | 0.009859 |
| **atlas.TOFusC-L** | atlas.sLOC-L | 4.690 | 0.009879 |
| **atlas.Hippocampus-R** | atlas.OFusG-L | 4.690 | 0.009879 |
| **atlas.iLOC-R** | atlas.sLOC-L | 4.680 | 0.009906 |
| **atlas.pTFusC-R** | atlas.OFusG-L | 4.680 | 0.009906 |
| **atlas.OFusG-R** | atlas.pPaHC-L | 4.660 | 0.010207 |
| **atlas.OFusG-L** | atlas.sLOC-L | 4.650 | 0.010275 |
| **atlas.OFusG-R** | atlas.pPaHC-R | 4.650 | 0.010275 |
| **atlas.TOFusC-L** | atlas.iLOC-L | 4.630 | 0.010535 |
| **atlas.iLOC-R** | atlas.pPaHC-R | 4.620 | 0.010535 |
| **atlas.OFusG-R** | atlas.iLOC-R | 4.610 | 0.01068 |
| **atlas.toITG-L** | atlas.Hippocampus-L | 4.600 | 0.010906 |
| **atlas.pPaHC-L** | atlas.OFusG-L | 4.600 | 0.010937 |
| **atlas.TOFusC-L** | atlas.pTFusC-L | 4.590 | 0.010937 |
| **atlas.pPaHC-L** | atlas.OFusG-R | 4.570 | 0.011228 |
| **atlas.toITG-L** | atlas.pPaHC-R | 4.550 | 0.011513 |
| **atlas.TOFusC-L** | atlas.pPaHC-L | 4.550 | 0.011611 |
| **atlas.aPaHC-L** | atlas.sLOC-L | 4.540 | 0.011669 |
| **atlas.iLOC-R** | atlas.pTFusC-R | 4.530 | 0.011911 |
| **atlas.pPaHC-L** | atlas.TOFusC-R | 4.510 | 0.012279 |
| **atlas.OFusG-L** | atlas.iLOC-L | 4.500 | 0.012439 |
| **atlas.pPaHC-R** | atlas.TOFusC-L | 4.470 | 0.013131 |
| **atlas.toITG-L** | atlas.sLOC-R | 4.440 | 0.013577 |
| **atlas.sLOC-L** | atlas.TOFusC-R | 4.410 | 0.014042 |
| **atlas.OFusG-R** | atlas.Hippocampus-L | 4.390 | 0.01418 |
| **atlas.toITG-R** | atlas.pTFusC-L | 4.350 | 0.014963 |
| **atlas.pPaHC-R** | atlas.iLOC-L | 4.270 | 0.016952 |
| **atlas.pTFusC-R** | atlas.sLOC-L | 4.160 | 0.01946 |
| **atlas.OFusG-R** | atlas.Hippocampus-R | 4.110 | 0.020818 |
| **atlas.Hippocampus-R** | atlas.OFusG-R | 4.100 | 0.020839 |
| **atlas.iLOC-L** | atlas.pPaHC-L | 4.100 | 0.020858 |
| **atlas.iLOC-R** | atlas.iLOC-L | 4.100 | 0.02094 |
| **atlas.iLOC-L** | atlas.toITG-R | 4.090 | 0.021119 |
| **atlas.TOFusC-R** | atlas.pPaHC-L | 4.090 | 0.021202 |
| **atlas.iLOC-L** | atlas.toITG-L | 4.050 | 0.021974 |
| **atlas.sLOC-L** | atlas.iLOC-R | 4.010 | 0.023359 |
| **atlas.TOFusC-L** | atlas.toITG-R | 4.000 | 0.023696 |
| **atlas.pPaHC-R** | atlas.iLOC-R | 3.980 | 0.0243 |
| **atlas.Hippocampus-R** | atlas.toITG-L | 3.960 | 0.024894 |
| **atlas.toITG-L** | atlas.aPaHC-L | 3.930 | 0.026051 |
| **atlas.iLOC-L** | atlas.sLOC-R | 3.920 | 0.026158 |
| **atlas.toITG-R** | atlas.pTFusC-R | 3.920 | 0.026297 |
| **atlas.pPaHC-L** | atlas.iLOC-L | 3.880 | 0.027906 |
| **atlas.aPaHC-L** | atlas.OFusG-R | 3.870 | 0.027953 |
| **atlas.OFusG-R** | atlas.aPaHC-L | 3.860 | 0.028146 |
| **atlas.pPaHC-R** | atlas.OFusG-R | 3.840 | 0.028785 |
| **atlas.Hippocampus-L** | atlas.toITG-R | 3.820 | 0.029911 |
| **atlas.toITG-L** | atlas.iLOC-R | 3.800 | 0.030496 |
| **atlas.sLOC-R** | atlas.toITG-L | 3.800 | 0.030508 |
| **atlas.sLOC-R** | atlas.TOFusC-R | 3.790 | 0.031037 |
| **atlas.iLOC-L** | atlas.pTFusC-L | 3.770 | 0.03192 |
| **atlas.pTFusC-L** | atlas.sLOC-R | 3.760 | 0.032237 |

| | | | |
|---|---|---|---|
| **atlas.OFusG-R** | atlas.pTFusC-R | 3.750 | 0.032441 |
| **atlas.iLOC-R** | atlas.OFusG-L | 3.730 | 0.033502 |
| **atlas.sLOC-R** | atlas.OFusG-L | 3.680 | 0.035796 |
| **atlas.TOFusC-R** | atlas.sLOC-R | 3.670 | 0.036142 |
| **atlas.toITG-L** | atlas.iLOC-L | 3.670 | 0.036151 |
| **atlas.pTFusC-L** | atlas.iLOC-R | 3.650 | 0.037216 |
| **atlas.pTFusC-L** | atlas.toITG-R | 3.640 | 0.038073 |
| **atlas.aPaHC-L** | atlas.OFusG-L | 3.630 | 0.038093 |
| **atlas.sLOC-L** | atlas.iLOC-L | 3.600 | 0.039544 |
| **atlas.iLOC-R** | atlas.toITG-L | 3.580 | 0.040615 |
| **atlas.toITG-R** | atlas.pPaHC-R | 3.550 | 0.041957 |
| **atlas.toITG-R** | atlas.Hippocampus-L | 3.540 | 0.042153 |
| **atlas.OFusG-L** | atlas.pTFusC-L | 3.520 | 0.042786 |
| **atlas.Hippocampus-L** | atlas.OFusG-L | 3.500 | 0.044317 |
| **atlas.Hippocampus-R** | atlas.iLOC-L | 3.470 | 0.046224 |
| **atlas.OFusG-L** | atlas.Hippocampus-L | 3.450 | 0.047012 |
| **atlas.sLOC-L** | atlas.toITG-R | 3.420 | 0.049055 |
| **atlas.pPaHC-L** | atlas.sLOC-L | 3.410 | 0.04959 |
| **atlas.sLOC-L** | atlas.TOFusC-L | 3.410 | 0.049791 |
